# MuDCoD: Multi-Subject Community Detection in Personalized Dynamic Gene Networks from Single Cell RNA Sequencing

**DOI:** 10.1101/2021.11.30.470619

**Authors:** Ali Osman Berk Şapcı, Shan Lu, Shuchen Yan, Ferhat Ay, Oznur Tastan, Sündüz Keleş

## Abstract

**Motivation:** With the wide availability of single-cell RNA-seq (scRNA-seq) technology, population-scale scRNA-seq datasets across multiple individuals and time points are emerging. While the initial investigations of these datasets tend to focus on standard analysis of clustering and differential expression, leveraging the power of scRNA-seq data at the personalized dynamic gene co-expression network level has the potential to unlock subject and/or time-specific network-level variation, which is critical for understanding phenotypic differences. Community detection from co-expression networks of multiple time points or conditions has been well-studied; however, none of the existing settings included networks from multiple subjects and multiple time points simultaneously. To address this, we develop MuDCoD for multi-subject community detection in personalized dynamic gene networks from scRNA-seq. MuDCoD builds on the spectral clustering framework and promotes information sharing among the networks of the subjects as well as networks at different time points. It clusters genes in the personalized dynamic gene networks and reveals gene communities that are variable or shared not only across time but also among subjects.

**Results:** Evaluation and benchmarking of MuDCoD against existing approaches reveal that MuDCoD effectively leverages apparent shared signals among networks of the subjects at individual time points, and performs robustly when there is no or little information sharing among the networks. Applications to population-scale scRNA-seq datasets of human-induced pluripotent stem cells during dopaminergic neuron differentiation and CD4+ T cell activation indicate that MuDCoD enables robust inference for identifying time-varying personalized gene modules. Our results illustrate how personalized dynamic community detection can aid in the exploration of subject-specific biological processes that vary across time.

**Availability:** MuDCoD is publicly available at https://github.com/bo1929/MuDCoD as a Python package. Implementation includes simulation and real-data experiments together with extensive documentation.

**Supplementary information:** Supplementary data are available at *Bioinformatics* online.

## 1 Introduction

The advent of single-cell RNA sequencing (scRNA-seq) has provided unparalleled insights into the transcriptional programs of different cell types and cellular stages at the individual cell level (Zeisel *et al*., 2015; Chen *et al*., 2017; Travaglini *et al*., 2020). Population-scale scRNA-seq datasets across multiple individuals and time points are becoming increasingly available (Mathys *et al*., 2019; HipSci Consortium *et al*., 2020, 2021; Soskic *et al*., 2022). These datasets make it possible to construct personalized, i.e., subject-specific, dynamic gene networks that vary across individuals and across time. Community structures emerge in these dynamic networks when there is strong local clustering of genes that are synchronized to function together. Unraveling the differentiation dynamics or deciphering pathways affected in various diseases requires discovering which genes cooperate in specific cellular processes. Thus, a key inference from the analysis of gene networks is detecting time/condition-varying gene modules that might correspond to genes cooperating in various biological processes. While these modules change over time due to the dynamic nature of the relations among the genes in the processes, they may also vary among different subjects. Variation and similarity among subjects at the network-level, when combined with subject-level phenotype information, can reveal critical information such as differential dynamics of the gene modules driving specific phenotypes. Thus, identifying gene modules by taking into account multiple subjects at once is of paramount importance.

Gene module detection methods for scRNA-seq data have mostly focused on the construction of static gene networks that capture a snapshot of time or a developmental epoch (Dai *et al*., 2019; Jackson *et al*., 2020). Many methods exist for detecting modules/communities in networks with or without specialization in multiple subjects or time points, both within and outside of genomics research (summarized in Table S1). These approaches leverage and combine stochastic block models (SBMs) (Matias and Miele, 2017a) with Kalman filters (Xu and Hero, 2014) or hidden Markov models (Ting *et al*., 2021), impose smoothing in spectral clustering (Chi *et al*., 2007; Liu *et al*., 2018), and utilize Steiner tree formulation (Norman and Cicek, 2019). Modularity maximization (Bassett *et al*., 2013; Betzel *et al*., 2019), non-negative matrix factorization (Ma *et al*., 2019), and network change point detection (NCPD) (Cribben and Yu, 2017) were also explored in community detection from brain and behavioral networks.

Recently, Liu *et al*. (2018) developed a global spectral clustering framework named PisCES for inferring communities in dynamic networks. PisCES detects gene modules at each time point by imposing module smoothness across time in a spectral clustering formulation. They show that such information sharing across the time domain improves community detection in dynamic networks (Liu *et al*., 2018). However, PisCES does not accommodate joint analysis of networks from multiple subjects, potentially overlooking information that could boost the signal for discovering communities. Here, we generalize the PisCES to detect communities in personalized dynamic gene networks and identify gene modules that vary or persist not only across time but also among subjects. Our method, named MuDCoD (**Mu**lti-subject **D**ynamic **Co**mmunity **D**etection), infers gene communities per subject and per time point by extending the temporal smoothness assumption to the subject dimension.

We evaluated MuDCoD with simulation experiments in a wide variety of scenarios. To this end, we extended the dynamic degree corrected block model (Dynamic DCBM) (Matias and Miele, 2017b), which provides a setting for time-varying networks to multi-subject scenarios. This multi-subject dynamic DCBM setting accommodates the characteristics of scRNA-seq-based personalized gene networks not only across time but also in the subject dimension. While there are several community detection methods within and outside of genomics, as we summarized above, they are limited in their applicability to handle dynamic multi-subject networks. After taking these approaches and the accessibility of their software into consideration, we compared MuDCoD with the recently developed multi-layer multi-subject modularity maximization technique (referred to as Betzel-2019 hereafter) (Betzel *et al*., 2019), PisCES, and static spectral clustering. We observed that MuDCoD markedly outperforms other methods when subjects share information at the network level and performs robustly otherwise. In addition to the simulation experiments, we also applied MuDCoD to scRNA-seq studies of long-term human induced pluripotent stem cells (iPSC) (HipSci Consortium *et al*., 2021) and activation of naive and memory human CD4+ T cells isolated from peripheral blood (Soskic *et al*., 2022). Our results highlight that MuDCoD is able to leverage information sharing across subjects and infer gene modules that contribute to network variation across subjects and time points. Downstream gene set enrichment analysis of inferred modules highlights persistent biological processes across subjects, as well as biological processes that are specific to subsets of subjects and/or time points. A detailed analysis of HipSci Consortium *et al*. (2021) dataset by leveraging the differentiation efficiency of subjects reveals that the gene modules of high differentiation efficiency subjects tend to exhibit higher variation across the differentiation time points compared to subjects with low differentiation efficiency. For Soskic *et al*. (2022) dataset, we observe that module structures inferred by MuDCoD and the dynamics of enriched biological process annotations for such modules are consistent with mechanisms governing CD4+ T cell activation.

## 2 Methods

### 2.1 Problem Formulation

We define a multi-subject dynamic gene co-expression network for discrete time steps *t* = 1, …, *T* and for subjects *s* = 1, …, *S* as a time series of undirected and unweighted graphs *𝒢* _*s*,1_, …, *𝒢*_*s,T*_ for each subject *s*. Given a multi-subject dynamic gene co-expression network, the key task is to infer the *community structure* for each time point and subject. The community structure of a gene co-expression network of size *G* is defined as a partition of genes into *K* disjoint cohesive modules/subsets, where *K* is a hyperparameter determining the number of communities. Each community is essentially a group of gene nodes, densely connected inside and loosely connected outside the community. The problem structure is shown in Fig. 1. In what follows, we use “community” and “module” interchangeably.

**Fig. 1.**
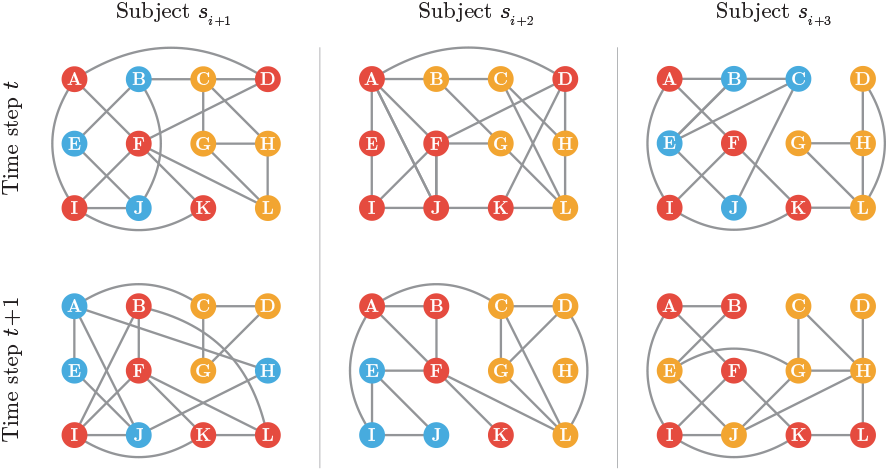
A schematic for multi-subject dynamic gene networks. A gene network is observed for each subject at each time step among a common set of nodes. The sets of edges vary among both the subjects and the time steps. These networks are estimated from scRNA-seq data and are expected to harbor communities that are conserved at varying levels among the subject and time dimensions. Different colors mark distinct communities, where the nodes (genes) within the same communities are depicted with the same color. MuDCoD assumes that communities change smoothly across both the subject and the time dimensions.

### 2.2 Promoting Signal Sharing Simultaneously in Subject and Time Dimensions

Let *A*_*s*,1_, …, *A*_*s,T*_ denote a time series of symmetric *G* × *G* adjacency matrices of networks *𝒢*_*s*,1_, …, *𝒢*_*s,T*_ varying across discrete time steps *t* = 1, …, *T*, for subjects *s* = 1, …, *S*. Let *L*_*s,t*_ denote the degree-normalized Laplacian of *A*_*s,t*_ as defined by:

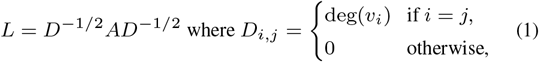

where deg(*υ*_*i*_) denotes the degree of node *υ*_*i*_.

Let *K* be fixed, and *V*_*s,t*_ ∈ ℝ^*G*×*K*^ denote a matrix with columns corresponding to the *K* leading eigenvectors of *L*_*s,t*_. A baseline strategy for inferring communities from these adjacency matrices is finding communities separately at each time step and for each individual. To detect communities that change longitudinally, PisCES (Liu *et al*., 2018) applies smoothing to the eigenvectors of *L*_*s,t*_ across time. We develop MuDCoD as a novel and empirically motivated (Supp. Section S5) extension of this framework. MuDCoD applies eigenvector smoothing across both the subject and the time dimensions to promote signal sharing across the subjects in addition to the time dimension. Let 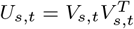 be the projection matrix onto the column space of *V*_*s,t*_ and define mean projection matrix

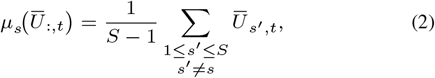

where *Ū*_*s,t*_ is the smoothed version of *U*_*s,t*_. We estimate *Ū*_*s,t*_ by:

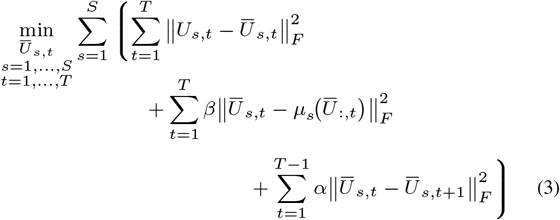

subject to *Ū*_*s,t*_, ∈ {*VV*^*T*^ : *V* ∈ ℝ^*G*×*K*^, *V*^*T*^*V* = *I* } ∀*s*, ∀*t*.

Here, the penalty term with parameter *α* enforces smoothness over the adjacent time points, whereas the term with parameter *β* constrains subject-specific variation from the mean time-dependent projection matrix *μ*_*s*_ (*Ū*_:,*t*_). Our formulation is based on the assumption that community structure of gene co-expression networks are expected to be similar across the subjects at a fixed time point (i.e., small 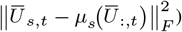) unless there is an explicit grouping variable (e.g., in a case and control setting). This assumption is empirically supported for the datasets we are considering in this paper (Supp. Section S5). The solution of Eq. 3 yields a series of smoothed mean projection matrices for each subject. Similar to PisCES, we propose to solve this non-convex optimization problem with the following iterative method:

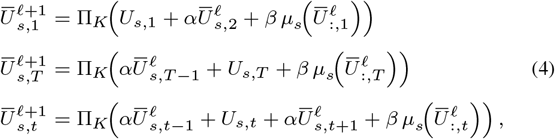

where *t* = 2, …, *T* − 1, *s* = 1, …, *S* for all iterations *ℓ* and the iterative algorithm is initialized by 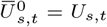 for 1 ≤ *t* ≤ *T, s* = 1, …, *S*. The mapping Π_*K*_(*M*) extracts the *K* leading eigenvectors, and is given by 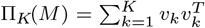, for a matrix *M*, where *υ*_1_, …, *υ*_*k*_ are the *K* leading eigenvectors of *M*. This formulation allows the model order *K* to be unknown and possibly varying over time by replacing Π_*K*_(*M*) with 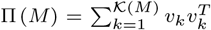. We provide a proof of convergence for this iterative algorithm and a range of hyperparameter values (α and β) for guaranteed convergence in Supp. Section S2.1. We utilize the eigengap statistics to select *𝒦*(*M*), and *α* and *β* are chosen with a re-sampling-based cross-validation strategy by Li *et al*. (2020), as we discuss in the next section. Supp. Fig. S1 illustrates the experimental convergence status for different *α* and *β* values.

### 2.3 Hyperparameter Selection

#### Selecting the number of communities

We allow the number of communities, *K*, to be unknown and possibly varying over time by exploiting the eigengap statistics. Shen and Cheng (2010) demonstrated that the degree-normalized Laplacian matrix and the correlation matrix significantly outperform the adjacency matrix, the standard Laplacian matrix, and the modularity matrix at identifying the community structure of networks. Therefore, we use eigenvalues of degree-normalized Laplacian matrix to estimate *K* as described in Supp. Section S1.2.

#### Cross-validation to tune *α* and *β*

We adopt the re-sampling based network cross-validation strategy of Li *et al*. (2020) to tune the hyperparameters *α* and *β*. This network cross-validation strategy makes sure to keep the node pairs in one fold when splitting the data into training and validation folds to avoid deleting edges and changing the network structure. Then, we apply grid search to find the best combination of hyperparameters *α* and *β*, where we define the best with respect to a higher DCBM likelihood function. In our experiments, we observed best performing *α* and *β* values to range between 0.25 and 0.75. Further details on this strategy are available in Supp. Section S1.1.

### 2.4 Prioritizing communities with CRank

Many community detection methods, including MuDCoD, fully partition network nodes into non-overlapping groups. However, in real-world datasets, only a few communities can typically be interpreted and linked to relevant underlying factors. Therefore, it is important to prioritize communities for downstream experimentation and further investigation.

CRank (Zitnik *et al*., 2018) prioritizes network communities by considering the structural features of each community and combining these features into an overall community score. Using metrics such as density, likelihood, allegiance, and boundary, CRank effectively ranks communities inferred by an external method.

We consider CRank as an optional but useful component of any analysis pipeline that includes MuDCoD. This is especially important for datasets with large numbers of subjects/time points, where aligning or matching subject/time-specific modules becomes infeasible. In our applications of MuDCoD, we benefitted from CRank by conducting a gene set enrichment analysis of only the prioritized communities for each subject.

## 3 Experimental Setup

### 3.1 Simulations

To evaluate MuDCoD, we extended the dynamic degree corrected block model (Dynamic-DCBM) (Xu and Hero, 2013) simulation set up to multi-subject setting (multi-subject dynamic degree corrected block model (MuS-Dynamic-DCBM)). This extension captures the overall characteristics of scRNA-seq-based personalized gene networks both in time and subject dimensions. In our computational experiments, we simulated modules from this model and assessed the performances of different methods in recovering the true modules.

The entries of the adjacency matrix *A*_*s,t*_ under MuS-Dynamic-DCBM are generated by the following Bernoulli distribution:

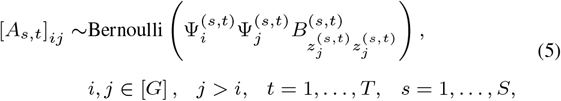

with [*A*_*s,t*_]_*ij*_ = [*A*_*s,t*_]_*ji*_, and [*A*_*s,t*_]_*ii*_ = 0. Here *z*^(*s,t*)^ ∈ [*K*]^*G*^ are vectors of community labels. More specifically, *i*-th entry 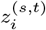 of *z*^(*s,t*)^denotes the module assignment of gene *i* for subject *s* at time point *t*. Ψ^(*s,t*)^ ∈ ℝ^*G*^are degree parameters and *B*^(*s,t*)^ ∈ [0, 1]^*K*×*K*^ is a connectivity matrix at time *t* for subject *s*. As we describe below with specific cases, variations in the generation process of 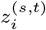 as a function of *s* and *t* provide a systematic way of modulating shared information, i.e., similarities, across the subject and time dimensions.

The degree parameters and the connectivity matrix (Ψ^(*s,t*)^ and *B*^(*s,t*)^) for subject *s* at time *t* are randomized as follows:

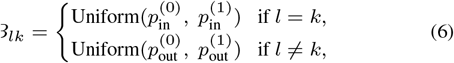

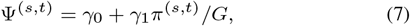

where 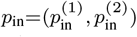 and 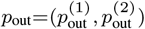 are in-cluster and between-cluster density parameters (*p*_in_ ≥ *p*_out_), respectively, *π* is a random permutation of 1:*G*, and *γ*_0_ and *γ*_1_ are determined based on the value of *G* and the desired degree distribution. We set *γ*_0_ = 0.5 and *γ*_1_ = 1 in our experiments. For the original network (*s*=1, *t*=1), community memberships of nodes are initialized from the following multinomial distribution: 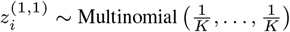.

Our extension MuS-Dynamic-DCBM spans two distinct multi-subject settings: Signal-Sharing-over-Subjects (SSoS) and Signal-Sharing-over-Time (SSoT) (Fig. 2). Both of these vary the degree of similarity between subject-specific networks while maintaining their time-dependent components and developing corresponding generation processes for community memberships across the time and subject dimensions. Signal-Sharing-over-Subjects (SSoS) enables modulation of signal sharing across subjects. For both settings, each subject’s network at time *t* is generated based on the corresponding module labels.

**Fig. 2.**
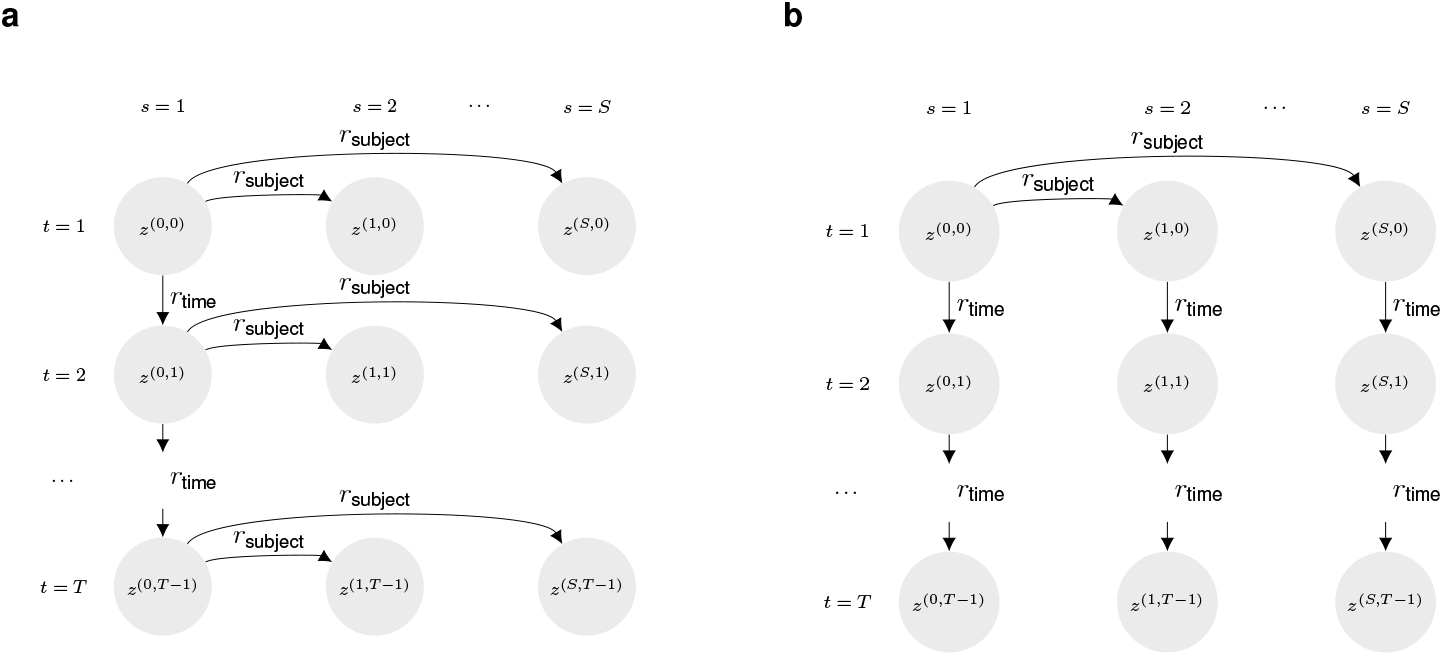
Multi-subject dynamic degree corrected block models (MuS-Dynamic-DCBM) for the two proposed settings. (a) SSoS setting: subjects evolve from a common ancestor at each time step *t*; and only the ancestor’s evolution over time is parameterized. (b) SSoT setting: subjects evolve from a common ancestor at *t* = 0; and then they evolve independently over time.

#### 3.1.1 Signal-Sharing-over-Subjects (SSoS)

In SSoS, gene modules of an “ancestor” progress over time with controllable similarity along the adjacent time points, and each subject’s set of gene modules at time *t* is a realization from this progressing ancestor (Fig. 2a). Let *s* = 1 index the subject for the ancestor network, and *z*^(1,*t*−1)^ denote its community labels for *t* = 2, …, *T*. In SSoS, *z*^(1,*t*−1)^progresses over time according to

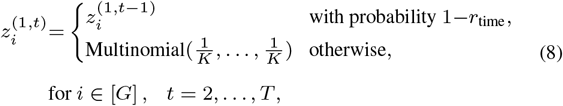

where *r*_time_ denotes the probability that a node *i* changes its previous community label, 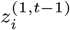. Then, community labels of other subjects for each time point *t* are generated as follows:

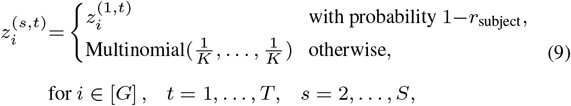

where *r*_subject_ is the probability that the node *i* changes community label while progressing from 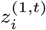.

#### 3.1.2 Signal-Sharing-over-Time (SSoT)

In contrast to SSoS, in SSoT, each subject’s gene modules start out from a common ancestor network at time *t* = 1 and progress over time independent of other subjects. In this setting, gene modules of a subject share similarities between time points (Fig. 2b). Let *s* = 1 index the subject for the ancestor network, and *z*^(1,1)^ denote its community labels at *t* = 1. The community labels of other subjects progress from the ancestor network only at time *t* = 1 according to:

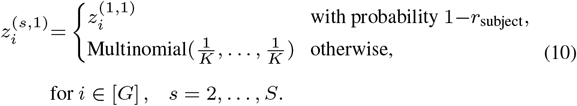

Then, each subject progresses independently over time by:

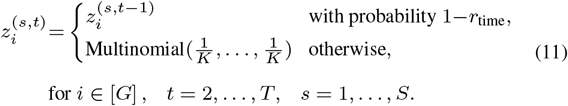

### 3.2 Experiments on Real Biological Datasets

We applied MuDCoD to two population-scale scRNA-seq datasets. In this section, we provide details on the data pre-processing steps of these applications. We remark that different data pre-processing steps were applied for the two datasets to accommodate differences in their overall levels of sparsity and the sequencing depths of the cells.

#### 3.2.1 scRNA-seq of iPS cells during dopaminergic neuron differentiation

The Jerber-2021 dataset (HipSci Consortium *et al*., 2021) harbors scRNA-seq data of 215 iPSC lines differentiating toward a mid-brain neural fate from the Human Induced Pluripotent Stem Cell Initiative (HipSci). Cells were collected for scRNA-seq profiling on days 11, 30, and 52 (day-11, day-30, and day-52). We considered the scRNA-seq count matrices from the three time points together for pre-processing. We retained genes expressed in at least 0.5% of the cells and selected top 3, 000 highly variable genes using the function `pp.filter_genes_dispersion`from Python package `Scanpy`. We normalized the count matrix for each donor using the `R` package `SCTransform`(Hafemeister and Satija, 2019) and adjusted for covariates “`pool_id`”, “`time_point`”, and “`treatment`” for batch correction. Leveraging the cell type annotations from the original publication (HipSci Consortium *et al*., 2021), we excluded day-11 from the analysis as it predominantly harbored progenitor cells which matured into young and mature neurons in the following time points. We focused on day-30 and day-52 both of which had dopaminergic neurons (DA), serotonin transporter (Sert), and ependymal-like 1 (Epen1) cell types. Furthermore, donors that (i) do not express the 3, 000 highly variable genes, i.e., have zero expression count across all cells, for one or more “time point-cell type” combinations; (ii) have fewer than 500 cells for any specific “time point-cell type” combination were filtered out for a robust estimation of gene co-expression networks. This resulted in 16, 22, and 8 donors, respectively, for DA, Sert, and Epen1 cell types. Finally, we constructed cell-type specific gene-gene Pearson correlation matrices for each donor at each time point.

#### 3.2.2 scRNA-seq of CD4+ T cells during activation

The Soskic-2022 dataset (Soskic *et al*., 2022) contains scRNA-seq profiling of 600k CD4+ T cells in a time course of activation after stimulation with anti-CD3/anti-CD28 beads. Cells were collected for scRNA-seq profiling before (0-hours) and at 16-hours, 40-hours, and 5-days after stimulation. We considered the scRNA-seq count matrices from the four time points together for pre-processing and retained genes expressed in at least 2% of the cells for initial gene filtering. All the ribosomal and mitochondrial genes were further removed. Datasets from specific time points and donor combinations with more than 200 cells were retained for the construction of networks. Due to the large sequencing depth differences between the cells profiled at different time points in this dataset, we applied `Dozer`(Lu and Keleş, 2023), which utilizes a Poisson measurement error model, to robustly estimate personalized gene co-expression networks. Furthermore, to improve the reliability and reproducibility of gene co-expression networks, we filtered out noisy and sparse genes using the “noise-ratio” metric from `Dozer` and retained 408 genes. Finally, this pre-processing yielded 79 healthy donors with co-expression networks of CD4+ Naive cells for 408 genes across the four time points.

For both datasets, the resulting co-expression matrices were then converted into unweighted adjacency matrices by keeping the top 5% of the edges (as commonly practiced in gene co-expression network construction from scRNA-seq (Iacono *et al*., 2019)) based on absolute values of their correlations. While this approach is commonly practiced and appears reasonable for these datasets (Supp. Section S8), alternatives that formally test for the edges in the co-expression networks could also be employed (Su *et al*., 2022).

## 4 Results

### 4.1 Performance Comparison with Simulation Experiments

We compared MuDCoD, PisCES, baseline static spectral clustering (“static”), and Betzel-2019 on simulated networks to assess their performance in community detection over the subject and time dimensions. We utilized the same selection procedure for the number of communities, *K*, when running PisCES, MuDCoD and “static” (Supp. Section S1.2). We used the network cross-validation approach to determine the tuning hyperparameters *α* and *β* for MuDCoD, and *α* for PisCES. Our choice of PisCES and “static” to compare MuDCoD with is motivated by PisCES’s overall established advantages over the two additional methods that Liu *et al*. (2018) had considered. Since Betzel-2019 has a different hyperparameter structure, we followed its default procedure, and provided the implementation details in Supp. Section S3. To measure the similarity between the inferred and the ground truth gene communities of each method, we used the adjusted Rand index (ARI) (Hubert and Arabie, 1985) as also used in Liu *et al*. (2018).

We conducted simulations under the SoSS and SSoT settings of MuS-Dynamic-DCBM with 16 subjects and for *T* ∈ {2, 4, 8}. The rest of the parameters were set as follows: network sizes *G*=500, number of communities *K*=10, in-cluster and out-cluster density parameters *p*_in_=(0.2, 0.4) and *p*_out_=(0.1, 0.1). We further considered varying *r*_time_ and *r*_subject_ levels. Here, *r*_time_ is the probability of a node changing its module label over two adjacent time points, and, similarly, *r*_subject_ is the probability of a node changing its module label across two subjects. For example, setting *r*_subject_=0 corresponds to the case where modules are similar across subjects, while *r*_subject_=0.5 corresponds to the setting where the modules of the subjects vary considerably and do not share any signal. Similarly, *r*_time_ = 0 and *r*_time_=0.5 yield constant and rapidly changing module labels across time, respectively. We conducted 100 simulation replicates and compared the methods based on the average ARI values over these replicates.

Fig. 3a and Fig. 3b summarize the results of these two general settings. For both settings, as *r*_subject_ and/or *r*_time_ increases, ARIs of MuDCoD, PisCES, and Betzel-2019 tend to decrease and, eventually, MuDCoD and PisCES perform similarly to the baseline “static”. This immediate observation is expected because, as the community structures among the subjects and/or along the adjacent time points become more dissimilar, promoting information sharing across either dimension is no longer advantageous. Compared to other methods, performance of Betzel-2019 appears to be most sensitive against increasing *r*_subject_ and *r*_time_. For *r*_subject_≥0.2 and/or *r*_time_≥0.2, Betzel-2019 achieves ARI values that are even less than (up to 0.2 less) those of the baseline static spectral clustering, and results in ARI values less than 0.2 when *r*_subject_=0.5 and/or *r*_time=_0.5.

**Fig. 3.**
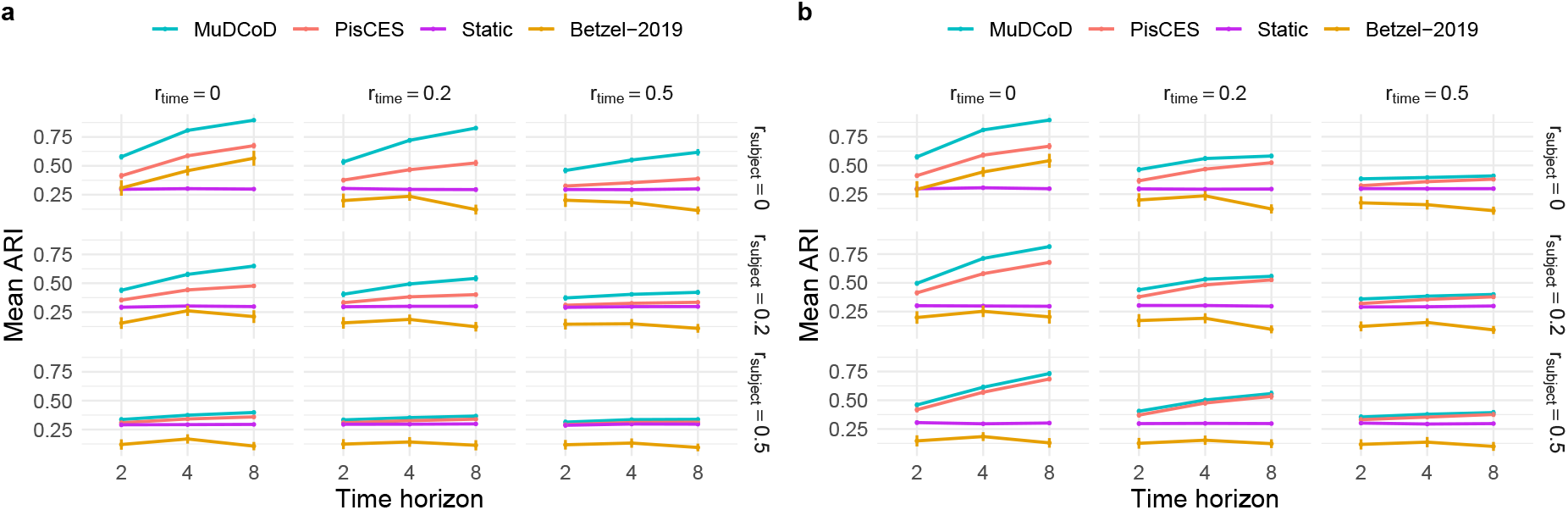
Evaluation of the identified communities under two different MuS-Dynamic-DCBM settings: (a) SSoS and (b) SSoT. The simulation parameters were set as follows: the network size is *G*=500, the number of class labels *K*=10, the in-cluster and out-cluster density parameters *p*_in_=(0.2, 0.4) and *p*_out_=(0.1, 0.1), number of subjects *S*=8, and the number of time points *T* ∈ {2, 4, 8}. *x*-axis is the number of time points *T*, and *y*-axis is the mean ARI of the inferred modules for all subjects and time steps across 100 simulation replicates.

For the SSoT setting, Fig. 3b reveals that even for the case with the most module heterogeneity along the subject and time dimensions (*r*_subject_ = 0.5 and *r*_time_ =0.5), PisCES and MuDCoD perform slightly better than both “static” and Betzel-2019. MuDCoD consistently performs better when there are varying levels of information sharing among the subjects (*r*_subject_ ∈ {0, 0.2, 0.5} and the modules are completely conserved across the time dimension (*r*_time_ =0.0), yielding more than 20% increase in the ARI with *T* = 8, compared to second best PisCES.

For the SSoS setting, Fig. 3a highlights MuDCoD as significantly outperforming other methods for relatively small *r*_subject_ values, and performing robustly against increasing *r*_time_ values. For the *r*_subject_ =0.5, MuDCoD performs no worse than other methods, if not negligibly better. When *T* =8, the average increase in ARI is about 20% and 8% for *r*_subject_ =0 and *r*_subject_ =0.2, respectively. The mean ARI increase is markedly high for some settings. For example, when *r*_subject_ = 0 and *r*_time_ =0.2, increase in the mean ARI reaches to almost 30% with *T* =8. Even with an extreme value of *r*_subject_ =0.5, which corresponds to high levels of heterogeneity between the subject modules, MuDCoD performs again at least as well as the other methods. PisCES and MuDCoD exhibit increasing performances as *T* increases, supporting the merits of information sharing across the time dimension. In contrast to PisCES and MuDCoD, Betzel-2019 appears to only benefit from increasing the time horizon when *r*_subject_ and *r*_subject_ are both set to zero.

### 4.2 Application of MuDCoD to discover multi-subject dynamic gene communities

#### 4.2.1 Analysis of HipSci Consortium *et al*. (2021) dataset

We next applied MuDCoD along with PisCES and static spectral clustering to discover personalized gene communities from the scRNA-seq data of human-induced pluripotent stem cells (HipSci Consortium *et al*., 2021). We specifically focused on cell types DA (16 donors, 1, 955 retained genes), Sert (22 donors and 2, 234 retained genes) and Epen1 (8 donors and 2, 475 retained genes) on days 30 and 52. Inferred communities varied in their sizes and densities (Table S2). Overall, the typical runtimes of MuDCoD and PisCES were comparable, ranging between 30 minutes to 1.5 hours for about 2, 000 genes (Supp. Section S7).

MuDCoD promotes the smoothness of spectral representations over donors within a time point; therefore, we would expect the gene modules across donors within a time point to be more similar to each other compared to PisCES and static spectral clustering, which treat each donor separately. We assessed to what extent this was realized in this application by calculating the normalized mutual information (NMI) scores (Strehl and Ghosh, 2003) among the discovered gene modules of the donors at each time point. Fig. 4a and Fig. 4b display the histograms of NMI scores between every pair of donors on day-30 and day-52. First, we observe that MuDCoD and PisCES infer gene modules that are more similar to each other across subjects, compared to static spectral clustering. Between MuDCoD and PisCES, MuDCoD tends to yield higher, but not statistically significant, NMI scores between donors on day-30 (Fig. 4a) and it leads to significantly higher scores at day-52 (Fig. 4b, Wilcoxon rank-sum test p-value of 0.0003), highlighting the impact of information sharing across the donors. Furthermore, this does not incur at the cost of additional smoothing over the time domain, as MuDCoD identified communities of donors display a level of heterogeneity comparable to those of PisCES across the time points (Fig. 4c).

**Fig. 4.**
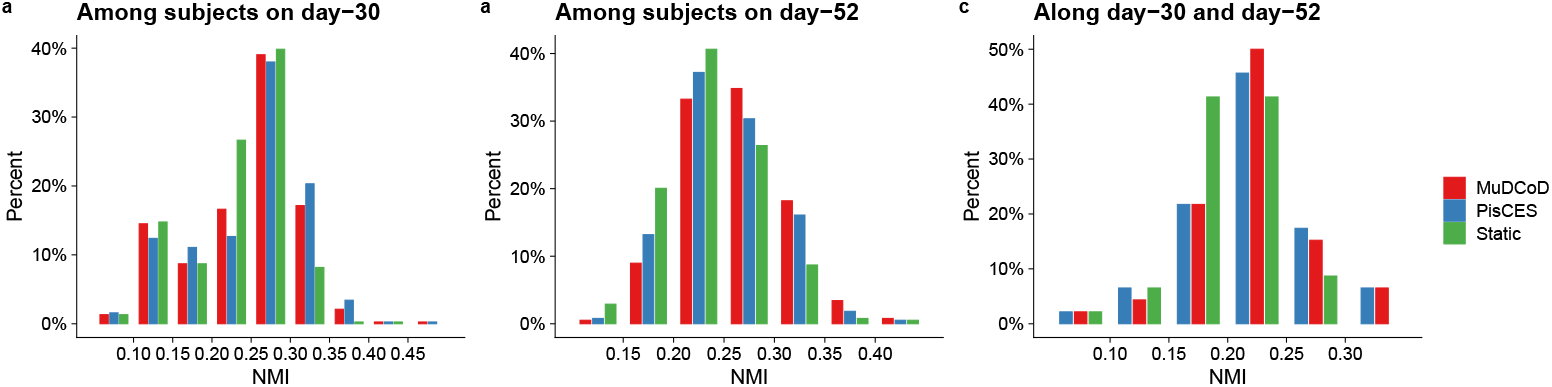
NMI scores between inferred gene modules of donor and time point pairs aggregated across all cell types. (a) and (b) quantify NMI scores between modules of every pair of donors on day-30 and day-52, respectively. (c) displays NMI scores between inferred gene modules of each donor on day-30 and on day-52. The *y*-axis denotes the percentage of donor pairs.

Following these general observations, we investigated the biological implications of MuDCoD results. To elucidate the biological processes that the modules might be involved in, we conducted gene set enrichment analysis of the discovered modules of each donor at each time point with `clusterProfiler` (Yu *et al*., 2012), with an emphasis on the Gene Ontology (GO) biological processes. We only focused on the communities with sizes larger than 20. Our analysis limited the background gene sets in `clusterProfiler` to only those genes that were used in the construction of the adjacency matrices and used the Benjamini-Hochberg procedure (Benjamini and Hochberg, 1995) for false discovery rate (FDR) control at 0.05. Supp. Fig. S3 highlights that some general biological processes, e.g., general cell-cycle and cell division-related processes, are enriched across the communities of all donors. Additionally, we also observe processes specific to subsets of donors, highlighting the personalized nature of the communities.

Next, to further explore the utility of our results, we leveraged the grouping of the donors based on the differentiation efficiency of their cells during dopaminergic neuron differentiation, as defined in HipSci Consortium *et al*. (2021). We specifically assessed whether donors with low and high differentiation efficiency exhibited varying levels of similarity among their modules inferred at different time points. Fig. 5a displays the NMI scores between the inferred modules of day-30 and day-52 for each donor, and suggests that donors with lower differentiation efficiency tend to have higher levels of similarity among their gene communities inferred at the two-time points (see Supp. Figs S8a and S9a for qualitatively similar results of PisCES and static spectral clustering). We then focused on cell type Epen1, which is not used in the definition of differentiation efficiency by (HipSci Consortium *et al*., 2021), and asked whether differences in the donor-specific communities are associated with the differentiation efficiency. This resulted in markedly higher module similarity within each group (i.e., high vs. low differentiation efficiency) and lower across groups (Fig. 5b). We observed similar trends with PisCES and static spectral clustering (Supp. Figs S8b and S9b). Furthermore, consistent with the differentiation dynamics (HipSci Consortium *et al*., 2021), we also observed relatively higher heterogeneity within the high group compared to the low group. Collectively, these observations support that personalized modules inferred by MuDCoD recapitulate the underlying differentiation efficiencies of donor iPS cells.

**Fig. 5.**
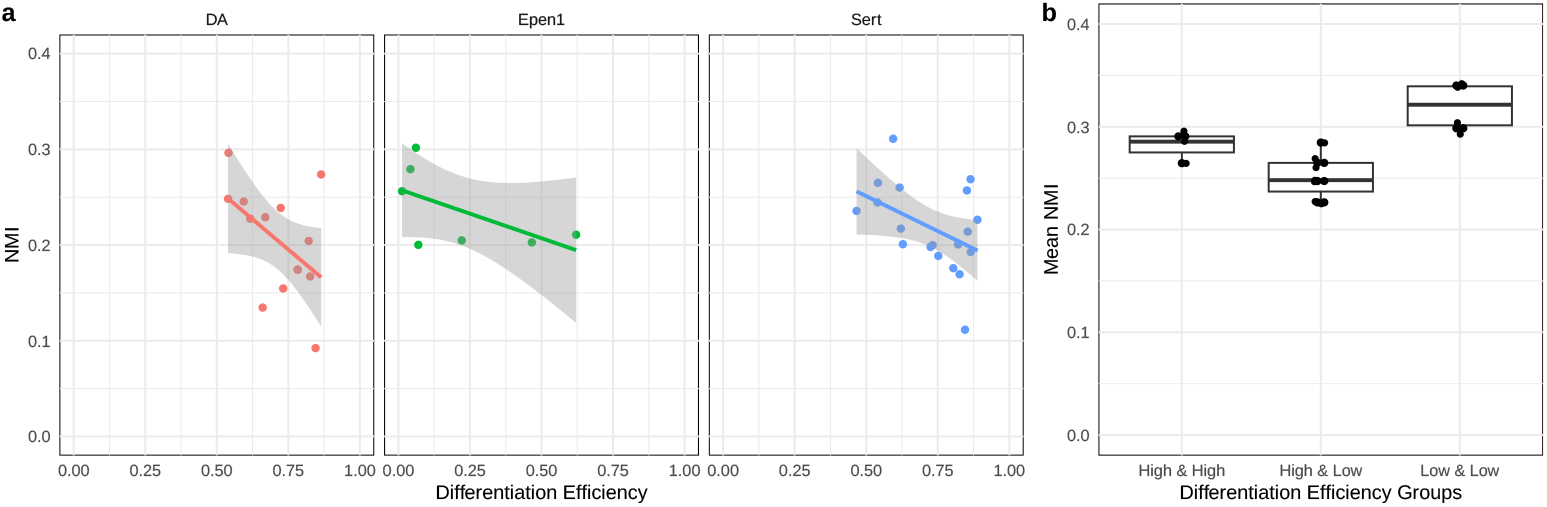
(a) NMI scores of each donor between the MuDCoD inferred modules on day-30 and on day-52 against the differentiation efficiency. For each cell type (DA, Sert, Epen1), each donor’s modules from day-30 and day-52 were compared with NMI and plotted against donor’s differentiation efficiency. (b) Comparison of the mean NMI scores within and between the low and high differentiation efficiency groups of Epen1 cells. Donor labels for differentiation efficiency were obtained from HipSci Consortium et al. (2021). NMI scores between pairs of donors are calculated based on their MuDCoD inferred modules. Differentiation efficiency groups were generated based on the percentiles of differentiation efficiency values across the donors, i.e., 50-th percentile (inclusive) corresponds to the low differentiation efficiency group.

#### 4.2.2 Analysis of Soskic *et al*. (2022) dataset

We applied MuDCoD along with PisCES and static spectral clustering to discover dynamic gene modules from the scRNA-seq data of CD4+ T cell activation (Soskic *et al*., 2022). This dataset includes both memory and naive CD4+ T cells isolated from peripheral blood. We focused on Naive CD4+ T cells, which have not yet encountered an antigen. The entire dataset captures transcriptional states of unstimulated cells (0-hours) and three time points (16-hours, 40-hours, and 5-days after the stimulation) of cell activation in 79 healthy donors, and 408 retained genes, as described in Section 3.2.2. Table S2 reports characteristics of inferred communitites at different time points. Similar to Jerber-2021, the typical runtimes of MuDCoD and PisCES in this dataset were comparable, usually ranging between 20 minutes to 1 hour (Supp. Section S7).

Similar to the analysis of HipSci Consortium *et al*. (2021) dataset, we first evaluated to what extent MuDCoD promoted smoothness of spectral representations among donors within a time point compared to PisCES by leveraging the NMI scores. Supp. Fig. S4 displays the histograms of NMI scores between all possible pairs of donors at time points 0-hours, 16-hours, 40-hours, and 5-days for modules inferred by MuDCoD, PisCES and static spectral clustering. Consistent with our expectation, MuDCoD tends to yield higher NMI scores (one-sided Wilcoxon rank-sum test *p*-values of 0.0003, 0.01 and 0.00019, respectively, for the time points 0-hours, 16-hours, and 5-days). The higher similarities among the gene modules across donors highlight the impact of information sharing across the donors at fixed time points.

Next, we assessed how the similarities between donors change as donor cells respond to stimulation over time. Specifically, we computed the mean NMI scores between each donor and the rest of the donors at all four time points separately. Figure 6a displays mean NMI scores that range between 0.05 and 0.2 across the time points. This comparison reveals that the inferred community structures of donors tend to be more similar at 16-hours after stimulation, as evidenced by the significant *p*-values (≤0.05) of the one-sided Wilcoxon rank sum tests comparing mean NMI scores at 16-hours with those at other time points. We note that 16-hours corresponds to the first measurement of gene expression right after stimulation of the T cells and before the first cell division. Hence, the increased similarity across donors may correspond to the concerted activity of T cell activation related genes and pathways. Analysis of PisCES and static spectral clustering output led to qualitatively similar results (Supp. Fig. S10A and S11A).

**Fig. 6.**
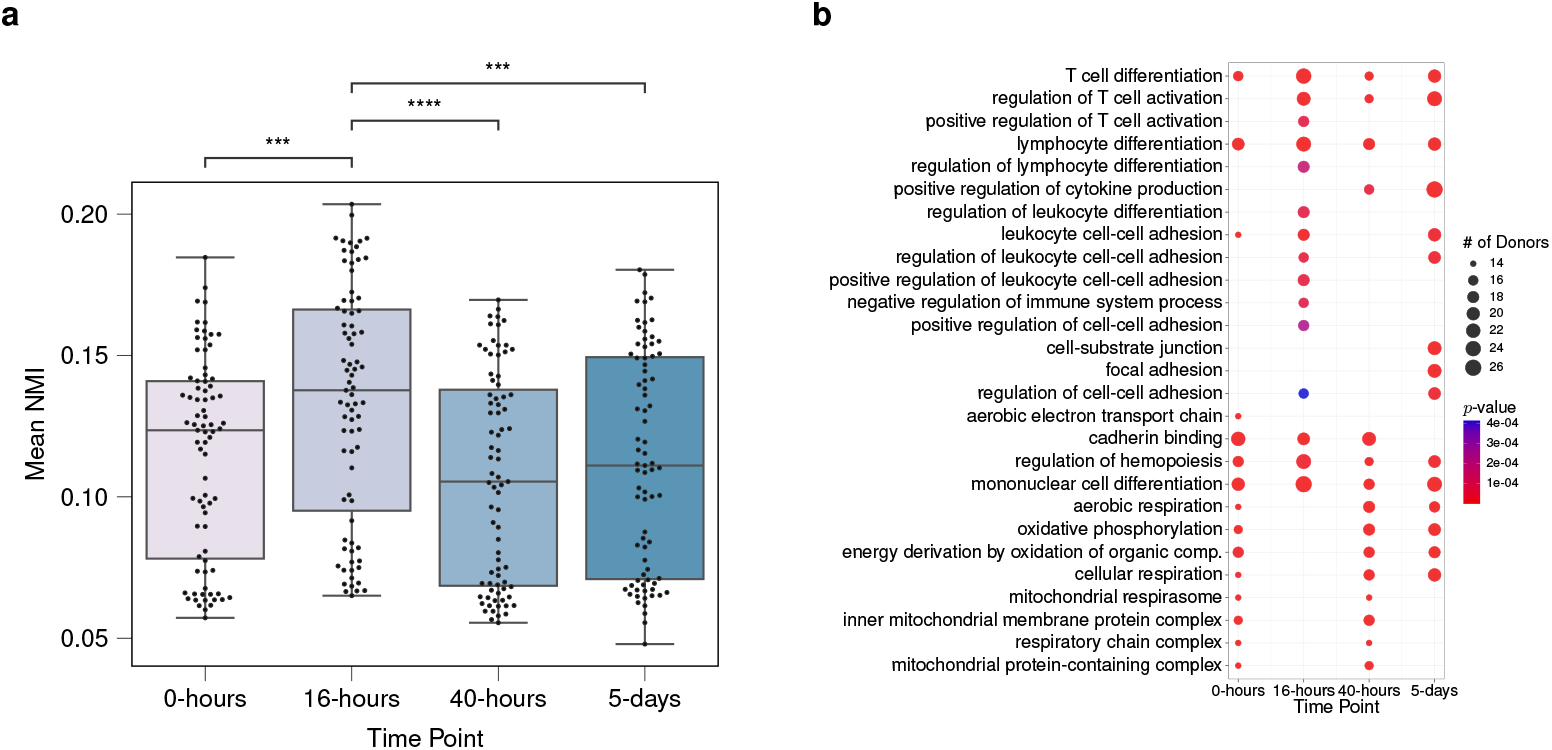
(a) Normalized mutual information scores between pairs of donors based on their gene modules inferred by MuDCoD at each time point. Each data point stands for a donor, and the *y*-axis denotes the mean of NMI scores between that donor and other donors at the corresponding time point. All the statistically significant comparisons with time point 16-hours from the one-sided Wilcoxon rank-sum test are marked (with *p*-value≤0.05), ∗∗∗ and ∗∗∗∗ stand for *p*-value≤0.001 and *p*-value≤0.0001, respectively. (b) Set of top fifteen frequent significantly enriched biological processes (with the adjusted *p*-values ≤0.05) of most prioritized communities contributed by each donor at different time points. Overall, 79 × 4=316 gene sets were contributed by 79 donors at 4 time points, and correspondingly, 316 separate enrichment analyses were performed. Displayed are significant biological processes, their corresponding number of appearances in the most prioritized communities of donors, and the minimum *p*-values of enrichment among those communities.

Following these general observations, we investigated the biological implications of MuDCoD results. MuDCoD fully partitions genes into modules. However, not all gene modules are necessarily interesting or biologically relevant. Furthermore, due to dropouts in scRNA-seq data, it is likely that only a small number of the inferred communities are biologically relevant (i.e., prioritized communities) and should be subject to further interpretation. We utilized CRank (Zitnik *et al*., 2018) to rank the resulting communities. Specifically, we ranked inferred communities of each donor’s network and identified the one with the highest rank, i.e., the most prioritized community. In the case of ties in CRank scores, we prioritized the larger community. This prioritization resulted in 79 × 4 = 316 gene sets when aggregated across donors and time points. Next, we conducted gene set enrichment analysis with GO biological processes for each of the prioritized modules using `clusterProfiler` (Yu *et al*., 2012). In order to summarize and elucidate the underlying biological processes at each time point, we calculated the frequencies of significant GO terms (FDR control at level 0.05 with the Benjamini-Hochberg procedure (Benjamini and Hochberg, 1995) at each time point). Figure 6b displays the top 15 most frequently enriched GO terms at each time point, along with their frequencies (i.e., number of appearances across donors) and minimum adjusted *p*-values across the donors (Supp. Fig. S10B and S11B for PisCES and statistc spectral clustering, respectively). Time point specific enriched biological processes appear consistent with the mechanisms governing CD4+ T cell activation (Hwang *et al*., 2020). For example, positive regulation of T cell activation is among the top 15 most frequent terms only at 16-hours and 40-hours after stimulation, and we start to observe positive regulation of cytokine production more frequently at 40-hours after stimulation. While these results require further analysis to reveal more detailed biological insights, MuDCoD successfully highlights how multi-subject module detection promoting module smoothness over the subject and time dimensions can yield gene modules that associate with specific biological conditions and phenotypes.

## 5 Conclusion

As advances in single-cell profiling technologies allow charting transcriptomes of individuals at unprecedented resolutions across developmental and cellular stages, analysis of personalized dynamic gene networks is poised to emerge as a critical tool for elucidating network-level variations among subjects to explain phenotypic variation. In this work, we developed a global spectral clustering framework that promotes information sharing among subjects and adjacent time points. MuDCoD utilizes co-expression networks to infer co-expression modules and can be effectively combined with any co-expression network construction method. For population-scale scRNA-seq data, we recommend using Dozer(Lu and Keleş, 2023), which employs a Poisson measurement error model to handle sparsity and correct gene pair correlation biases. In addition, thresholding the subject-specific co-expression networks to obtain adjacency matrices should also be tailored by taking into account the gene-gene correlation distribution characteristics of the datasets. Recent developments in testing of gene-gene correlations, like CS-CORE (Su *et al*., 2022), offer potential insights into binarizing co-expression networks.

Our simulation experiments highlighted the superior performance of MuDCoD over existing alternatives. Applications to two population-scale scRNA-seq datasets revealed that MuDCoD infers biologically meaningful and relevant communities in multi-subject dynamic scRNA-seq datasets. While our current framework encourages information sharing among all subjects, it can be extended to accommodate groupings among subjects where the communities are persistent only within the group members. Such an extension could be useful, especially when an apparent grouping such as healthy versus diseased subjects exists in the dataset. Finally, we expect this multi-subject setting to be useful in other domains to infer network community structures with the availability of multiple samples.

## References

Bassett, D. S. et al. (2013). Robust detection of dynamic community structure in networks. Chaos: An Interdisciplinary Journal of Nonlinear Science, 23(1), 013142.

Benjamini, Y. and Hochberg, Y. (1995). Controlling the False Discovery Rate: A Practical and Powerful Approach to Multiple Testing. Journal of the Royal Statistical Society. Series B (Methodological), 57(1), 289–300.

Betzel, R. F. et al. (2019). The community structure of functional brain networks exhibits scale-specific patterns of inter- and intra-subject variability. NeuroImage, 202, 115990.

Chen, Y.-J. J. et al. (2017). Single-cell RNA sequencing identifies distinct mouse medial ganglionic eminence cell types. Scientific Reports, 7(1), 1–11.

Chi, Y. et al. (2007). Evolutionary spectral clustering by incorporating temporal smoothness. In Proceedings of the 13th ACM SIGKDD international conference on Knowledge discovery and data mining, pages 153–162.

Cribben, I. and Yu, Y. (2017). Estimating whole-brain dynamics by using spectral clustering. Journal of the Royal Statistical Society: Series C (Applied Statistics), 66(3), 607–627.

Dai, H. et al. (2019). Cell-specific network constructed by single-cell RNA sequencing data. Nucleic Acids Research, 47(11), e62–e62.

Hafemeister, C. and Satija, R. (2019). Normalization and variance stabilization of single-cell RNA-seq data using regularized negative binomial regression. Genome biology, 20(1), 1–15.

HipSci Consortium et al. (2020). Single-cell RNA-sequencing of differentiating iPS cells reveals dynamic genetic effects on gene expression. Nature Communications, 11(1), 810.

HipSci Consortium et al. (2021). Population-scale single-cell RNA-seq profiling across dopaminergic neuron differentiation. Nature Genetics, 53(3), 304–312.

Hubert, L. and Arabie, P. (1985). Comparing partitions. Journal of classification, 2(1), 193–218.

Hwang, J.-R. et al. (2020). Recent insights of T cell receptor-mediated signaling pathways for T cell activation and development. Experimental & Molecular Medicine, 52(5), 750–761.

Iacono, G. et al. (2019). Single-cell transcriptomics unveils gene regulatory network plasticity. Genome biology, 20(1), 1–20.

Jackson, C. A. et al. (2020). Gene regulatory network reconstruction using single-cell RNA sequencing of barcoded genotypes in diverse environments. eLife, 9, e51254.

Li, T. et al. (2020). Network cross-validation by edge sampling. Biometrika, 107(2), 257–276.

Liu, F. et al. (2018). Global spectral clustering in dynamic networks. Proceedings of the National Academy of Sciences, 115(5), 927–932.

Lu, S. and Keleş, S. (2023). Debiased personalized gene coexpression networks for population-scale scrna-seq data. Genome Research, page gr277363. https://genome.cshlp.org/content/early/2023/07/11/gr.277363.122.

Ma, X. et al. (2019). Detecting evolving communities in dynamic networks using graph regularized evolutionary nonnegative matrix factorization. Physica A: Statistical Mechanics and its Applications, 530, 121279.

Mathys, H. et al. (2019). Single-cell transcriptomic analysis of Alzheimer’s disease. Nature, 570(7761), 332–337.

Matias, C. and Miele, V. (2017a). Statistical clustering of temporal networks through a dynamic stochastic block model. Journal of the Royal Statistical Society: Series B, 79(4), 1119–1141.

Matias, C. and Miele, V. (2017b). Statistical clustering of temporal networks through a dynamic stochastic block model. Journal of the Royal Statistical Society: Series B, 79(4), 1119–1141.

Norman, U. and Cicek, A. E. (2019). ST-Steiner: a spatio-temporal gene discovery algorithm. Bioinformatics, 35(18), 3433–3440.

Shen, H.-W. and Cheng, X.-Q. (2010). Spectral methods for the detection of network community structure: a comparative analysis. Journal of Statistical Mechanics: Theory and Experiment, 2010(10), P10020.

Soskic, B. et al. (2022). Immune disease risk variants regulate gene expression dynamics during CD4+ T cell activation. Nature Genetics, 54(6), 817–826. Number: 6 Publisher: Nature Publishing Group.

Strehl, A. and Ghosh, J. (2003). Cluster Ensembles – a Knowledge Reuse Framework for Combining Multiple Partitions. Journal of Machine Learning Research, 3, 583–617.

Su, C. et al. (2022). Cell-type-specific co-expression inference from single cell rna-sequencing data. bioRxiv, pages 2022–12.

Ting, C.-M. et al. (2021). Detecting Dynamic Community Structure in Functional Brain Networks Across Individuals: A Multilayer Approach. IEEE Transactions on Medical Imaging, 40(2), 468–480. Conference Name: IEEE Transactions on Medical Imaging.

Travaglini, K. J. et al. (2020). A molecular cell atlas of the human lung from single-cell RNA sequencing. Nature, 587(7835), 619–625.

Xu, K. S. and Hero, A. O. (2013). Dynamic Stochastic Blockmodels: Statistical Models for Time-Evolving Networks. In A. M. Greenberg, W. G. Kennedy, and N. D. Bos, editors, Social Computing, Behavioral-Cultural Modeling and Prediction, Lecture Notes in Computer Science, pages 201–210, Berlin, Heidelberg. Springer.

Xu, K. S. and Hero, A. O. (2014). Dynamic stochastic blockmodels for time-evolving social networks. IEEE Journal of Selected Topics in Signal Processing, 8(4), 552–562.

Yu, G. et al. (2012). clusterProfiler: an R Package for Comparing Biological Themes Among Gene Clusters. OMICS: A Journal of Integrative Biology, 16(5), 284–287.

Zeisel, A. et al. (2015). Cell types in the mouse cortex and hippocampus revealed by single-cell RNA-seq. Science, 347(6226), 1138–1142.

Zitnik, M. et al. (2018). Prioritizing network communities. Nature Communications, 9(1), 2544. Number: 1 Publisher: Nature Publishing Group.

