## Supplementary for "MuDCoD: Multi-Subject Community Detection in Personalized Dynamic Gene Networks from Single Cell RNA Sequencing"

### S1 Hyperparameter Selection for MuDCoD

#### S1.1 Cross-validation to Tune Hyperparameters $\alpha$ and $\beta$

We implemented a cross-validation scheme based on Li *et al.* [8] to select the tuning parameters $\alpha$ and $\beta$ as follows.

1. Randomly split the $\binom{G}{2} \cdot T \cdot S$ dyads of $A_{1,1}, \ldots, A_{S,T}$ into $F$ folds. Let $Q$ denote the the set of candidate pairs for $(\alpha, \beta)$.

2. For each fold $f = 1, \ldots, F$ and time-subject pair $(s, t)$, define $A_{s,t}^{(f)} \in \mathbb{R}^{G \times G}$ as:

$$\left[A_{s,t}^{(f)}\right]_{ij} = [A_{s,t}]_{ij} \, \mathbf{I}\left\{(i, j, s, t) \in f\right\}, \tag{1}$$

where $\mathbf{I}$ denotes the indicator function.

3. Apply a low-rank matrix completion algorithm [8] to $A_{s,t}^{(f)}$ to impute the zeroed entries, and let $\hat{A}_{s,t}^{(f)}$ denote the resulting matrix. Let $\overline{U}_{1,1}^{(\alpha,\beta,f)}, \ldots, \overline{U}_{S,T}^{(\alpha,\beta,f)}$ denote MuDCoD's output with smoothing parameters $\alpha$, $\beta$ and imputed $\hat{A}_{1,1}^{(f)}, \ldots, \hat{A}_{S,T}^{(f)}$.

4. Evaluate $\mathcal{L}^{(\alpha,\beta,f)}$, the log likelihood of the fitted DCBM on the $f$-th fold:

$$\mathcal{L}^{(\alpha,\beta,f)} = \sum_{(i,j,s,t) \in f} [A_{s,t}]_{ij} \log(\left[P_{(s,t)}\right]_{ij}) + (1 - [A_{s,t}]_{ij}) \log(1 - \left[P_{(s,t)}\right]_{ij}), \tag{2}$$

where $\left[P_{(s,t)}\right]_{ij} \equiv \left[P_{(s,t)}^{(f)}\right]_{ij}$ is the estimated probability of connection between node $i$ and node $j$ at time $t$ for subject $s$ under the fitted DCBM, and is given by:

$$\left[P_{(s,t)}\right]_{ij} = d_i^{(s,t)} d_j^{(s,t)} B^{(s,t)}(\hat{z}_i^{(s,t)}, \hat{z}_i^{(s,t)}). \tag{3}$$

Here, $\hat{z}_i^{(s,t)} \equiv \hat{z}_i^{(s,t,f)}$ is the estimated community node $i$ at time $t$ as given by clustering on the eigenvectors of $\overline{U}_{s,t}^{(\alpha,\beta,f)}$ with $\mathcal{K}(\overline{U}_{s,t}^{(\alpha,\beta,f)})$ classes where $d_i^{(s,t)} \equiv d_i^{(s,t,f)}$ is the estimated DCBM degree parameter for node $i$, and is given by:

$$d_i^{(s,t)} = \sum_j \left[\hat{A}_{s,t}^{(f)}\right]_{ij}, \tag{4}$$

and $B^{(s,t)} \equiv B^{(s,t,f)} \in \mathbb{R}^{K \times K}$ is the estimated DCBM density parameter matrix, and is given by:

$$\left[B^{(s,t)}\right]_{kl} = \left(\sum_{(i,j)} \left[\hat{A}_{s,t}^{(f)}\right]_{ij} \mathbf{I}\left\{\hat{z}_i^{(s,t)} = k,\ \hat{z}_j^{(s,t)} = l\right\}\right) / \left(\sum_{(i,j)} d_i^{(s,t)} d_j^{(s,t)} \mathbf{I}\left\{\hat{z}_i^{(s,t)} = k,\ \hat{z}_j^{(s,t)} = l\right\}\right). \tag{5}$$

5. Let $\mathcal{L}^{(\alpha,\beta)} = \frac{1}{F}\sum_f \mathcal{L}^{(\alpha,\beta,f)}$ and return the tuning parameter pair $(\alpha, \beta) \in Q$ giving the best value for $\mathcal{L}^{(\alpha,\beta)}$.

#### S1.2 Selecting the Number of Communities $\mathcal{K}$

We allow the number of communities, $K$, to be unknown and possibly varying over time by exploiting the eigengap statistics. Results of Shen and Cheng [15] demonstrate that the degree-normalized Laplacian matrix and the correlation matrix significantly outperform the adjacency matrix, the standard Laplacian matrix, and the modularity matrix at identifying the community structure of networks. Therefore, we consider eigenvalues of degree-normalized Laplacian matrix $L$ to detect $K$.

Given $L \in \mathbb{R}^{G \times G}$ with eigenvalues $|\lambda_1|, \geq \cdots \geq |\lambda_G|$, define $\mathcal{K} : \mathbb{R}^{G \times G} \mapsto \mathbb{N}$ as a function that computes the number of modules, i.e., the number of eigenvectors, as:

$$\mathcal{K}(M) = \min\left\{K \ : \ |\lambda_i| - |\lambda_{(i+1)}| < \delta \text{ for all } i > K\right\}. \tag{6}$$

Here, $\delta$ denotes the threshold for the "noise" eigenvalues. Let $v_1, \ldots, v_{\mathcal{K}(M)}$ denote the eigenvectors of $M$ corresponding to the $\mathcal{K}(M)$ largest eigenvalues in absolute value. Then, $\Pi_K(M)$ can be replaced with $\Pi(M)$, which is defined as:

$$\Pi(M) = \sum_{k=1}^{\mathcal{K}(M)} v_k v_k^T. \tag{7}$$

Based on the assumption of independent observation noise across dyads (i.e., node/gene pairs), Liu *et al.* [9] simulates Erdős-Renyi normalized Laplacian matrices $L^{(ER)}$ with sizes and densities matching to observed $L$ to determine the threshold $\delta$ for the corresponding networks generated with DCBM. We adapt the same approach and compute $\delta$ by

$$\delta = \text{quantile}_{0.95}\left[max\left\{|\lambda_i^{(ER)}| - |\lambda_{i+1}^{(ER)}|,\ i \geq 2\right\}\right]. \tag{8}$$

We also note that the assumption of dyadically independent observation noise may not be appropriate for the real scRNA-seq co-expression networks. To accommodate this possibility, we compute the threshold $\delta$ as the largest eigengap excluding $\lambda_1$:

$$\delta = \max\left\{|\lambda_i| - |\lambda_{(i+1)}|,\ i \geq\right\}. \tag{9}$$

Finally, $\mathcal{K}$ is computed using the threshold values $\delta$ obtained with both approaches described above, and we pick whichever results in more communities.

### S2 Convergence of the MuDCoD's Iterative Algorithm

#### S2.1 Theoretical Convergence

**Definition** (Estimated Mean Projection Matrix)**.** Mean projection matrix estimation at time step $t$, for subject $s$, is given by averaging all other subjects as follows:

$$\mu_s\left(\bar{U}_{:,t}\right) = \frac{1}{S-1} \sum_{\substack{1 \leq s' \leq S \\ s' \neq s}} \bar{U}_{s',t}.$$

**Definition** (Contraction Mapping)**.** Let $\mathcal{X}$ be a metric space with associated distance metric d. Then a mapping $\mathrm{F} : \mathcal{X} \mapsto \mathcal{X}$ is called contraction mapping if it satisfies for some $0 \leq \gamma < 1$:

$$\mathrm{d}(\mathrm{F}(x), \mathrm{F}(y)) \leq \gamma \, \mathrm{d}(x, y).$$

**Theorem 1** (Contraction Mapping Theorem)**.** *Let* $\mathrm{F} : \mathcal{X} \mapsto \mathcal{X}$ *be a contraction mapping. The* F *has a unique fixed point* $x^*$ *satisfying* $\mathrm{F}(x^*) = x^*$*. Furthermore, the sequence* $x_{n+1} = \mathrm{F}\, x_n$ *(with* $x_0$ *arbitrarily initialized) satisfies* $x_n \to x^*$*.*

Eigenvector smoothing iterations can be written in terms of an operator $\mathrm{G} = (\mathrm{G}_{1,1}, \ldots, \mathrm{G}_{S,1}, \cdots, \mathrm{G}_{1,T}, \ldots, \mathrm{G}_{S,T})$, given by:

$$\begin{aligned}
\mathrm{G}_{s,1}\left(\bar{U}_{1,1}, \ldots, \bar{U}_{S,1}, \cdots, \bar{U}_{1,T}, \ldots, \bar{U}_{S,T}\right) &= \Pi_K\left(U_{s,1} + \alpha \bar{U}_{s,2} + \beta \mu_s\left(\bar{U}_{:,1}\right)\right) \\
\mathrm{G}_{s,t}\left(\bar{U}_{1,1}, \ldots, \bar{U}_{S,1}, \cdots, \bar{U}_{1,T}, \ldots, \bar{U}_{S,T}\right) &= \Pi_K\left(U_{s,t} + \alpha \bar{U}_{s,t+1} + \alpha \bar{U}_{s,t-1} + \beta \mu_s\left(\bar{U}_{:,t}\right)\right) \\
\mathrm{G}_{s,T}\left(\bar{U}_{1,1}, \ldots, \bar{U}_{S,1}, \cdots, \bar{U}_{1,T}, \ldots, \bar{U}_{S,T}\right) &= \Pi_K\left(U_{s,T} + \alpha \bar{U}_{s,T-1} + \beta \mu_s\left(\bar{U}_{:,T}\right)\right)
\end{aligned}$$

so that we can write $\bar{U}^{\ell+1}_{1:S,1:T} = \mathrm{G}\left(\bar{U}^{\ell}_{1:S,1:T}\right)$, where $\bar{U}_{1:S,1:T} \equiv \left(\bar{U}_{1,1}, \ldots, \bar{U}_{S,1}, \cdots, \bar{U}_{1,T}, \ldots, \bar{U}_{S,T}\right)$.

**Theorem 2** (Davis and Kahan Theorem)**.** *Let* $\Sigma$ *and* $\Sigma'$ *be symmetric, suppose* $S \subset \mathbb{R}$ *is an interval, and suppose for some positive integer* $K$ *that* $V, V'$ *in*$\mathbb{R}^{n \times K}$*, and the columns of* $V (V')$ *form an orthonormal basis for the sum of eigenspaces of* $\Sigma(\Sigma')$ *associated with the eigenvalues of* $\Sigma(\Sigma')$ *in* $S$*. Let* $\delta$ *be the minimum of distance between any eigenvalues of* $\Sigma$ *in* $S$ *between any eigenvalues of* $\Sigma$ *not in* $S$*. Then, there exists an orthogonal matrix* $R \in \mathbb{R}^{K \times K}$ *such that:*

$$\|VR - V'\|_F \leq \frac{\sqrt{2}}{\delta} \|\Sigma - \Sigma'\|_F. \tag{10}$$

**Lemma 1.** *A necessary condition for* $\bar{U}^*_{1:S,1:T}$ *to be a global minimum of the optimization problem given in Eq. 51 is that* $\bar{U}^*_{1:S,1:T} = \mathrm{G}(\bar{U}^*_{1:S,1:T})$*.*

*Proof.* Let $\mathcal{V} = \left\{ V \in \mathbb{R}^{n \times K} : V^T V = I \right\}$, so that $\mathcal{U} = \left\{ VV^T : V \in \mathcal{V} \right\}$ denotes the feasible region of each $\bar{U}_{st}$ in the optimization problem. A necessary condition for $\bar{U}^*_{1:S,1:T}$ to be the global minimum of the optimization problem is that each $\bar{U}^*_{st}$ for must minimize the objective when the other variables are held constant:

$$\bar{U}^*_{s,1} = \underset{\bar{U}_{s,1} \in \mathcal{U}}{\arg\min} \left\|U_{s,1} - \bar{U}_{s,1}\right\|_F^2 + \alpha \left\|\bar{U}_{s,1} - \bar{U}^*_{s,2}\right\|_F^2 + \beta \left\|\bar{U}_{s,1} - \mu_s(\bar{U}^*_{:,1})\right\|_F^2 \tag{11}$$

$$\bar{U}^*_{s,t} = \underset{\bar{U}_{s,t} \in \mathcal{U}}{\arg\min} \left\|U_{s,t} - \bar{U}_{s,t}\right\|_F^2 + \alpha \left\|\bar{U}_{s,t} - \bar{U}^*_{s,t+1}\right\|_F^2 + \alpha \left\|\bar{U}_{s,t} - \bar{U}^*_{s,t-1}\right\|_F^2 + \beta \left\|\bar{U}_{s,t} - \mu_s(\bar{U}^*_{:,t})\right\|_F^2 \tag{12}$$

$$\bar{U}^*_{s,T} = \underset{\bar{U}_{s,T} \in \mathcal{U}}{\arg\min} \left\|U_{s,T} - \bar{U}_{s,T}\right\|_F^2 + \alpha \left\|\bar{U}_{s,T} - \bar{U}^*_{s,T-1}\right\|_F^2 + \beta \left\|\bar{U}_{s,T} - \mu_s(\bar{U}^*_{:,T})\right\|_F^2. \tag{13}$$

To prove the lemma, it thus suffices to show this condition is equivalent to $\bar{U}^*_{s,t} = \mathrm{G}_{s,t}(\bar{U}^*_{1:S,1:T})$ for each $s$ and $t$. Using the identity $\|M\|_F^2 = \mathrm{Tr}\left(M^T M\right)$ and the fact that $\|U\|_F^2 = K$ for all $U \in \mathcal{U}$, algebraic manipulation of Eq. 12 yields

$$\begin{aligned}
\bar{U}^*_{s,t} &= \underset{\bar{U}_{s,t} \in \mathcal{U}}{\arg\max}\ \mathrm{Tr}\left(\bar{U}^T_{s,t} U_{s,t}\right) + \alpha\, \mathrm{Tr}\left(\bar{U}^T_{s,t} \bar{U}^*_{s,t+1}\right) + \alpha\, \mathrm{Tr}\left(\bar{U}^T_{s,t} \bar{U}^*_{s,t-1}\right) + \beta\, \mathrm{Tr}\left(\bar{U}^T_{s,t} \mu_s(\bar{U}^*_{:,t})\right) && (14) \\
&= \underset{\bar{U}_{s,t} \in \mathcal{U}}{\arg\max}\ \mathrm{Tr}\left(\bar{U}^T_{s,t}\left(U_{s,t} + \alpha \bar{U}^*_{s,t+1} + \alpha \bar{U}^*_{s,t-1} + \beta \mu_s(\bar{U}^*_{:,t})\right)\right) && (15)
\end{aligned}$$

Since $\overline{U}^*_{s,t} \in \mathcal{U}$, we can let $\overline{U}^*_{s,t} = \overline{V}^* \overline{V}^{*T}$ for some $\overline{V} \in \mathcal{V}$, so that Eq. 14 implies that

$$\overline{V}^* = \underset{\overline{V} \in \mathcal{V}}{\arg\max} \ \mathrm{Tr}\left(\overline{V}^T \left(U_{s,t} + \alpha \overline{U}^*_{s,t+1} + \alpha \overline{U}^*_{s,t-1} + \beta \mu_s(\overline{U}^*_{:,t})\right) \overline{V}\right) \tag{16}$$

$$= \mathrm{Eigvec}_K \left(U_{s,t} + \alpha \overline{U}^*_{s,t+1} + \alpha \overline{U}^*_{s,t-1} + \beta \mu_s(\overline{U}^*_{:,t})\right) \tag{17}$$

where $\mathrm{Eigvec}_K$ denotes the $n \times K$ matrix in $\mathcal{V}$ whose columns are the first $K$ eigenvectors. Here we have used the fact that $U_{s,t}, \overline{U}^*_{s,t+1}, \overline{U}^*_{s,t-1}$ and $\mu_s(\overline{U}^*_{:,t})^\dagger$p are positive semi-definite matrices.

$$\overline{U}^*_{s,t} = \Pi_K \left(U_{s,t} + \alpha \overline{U}^*_{s,t+1} + \alpha \overline{U}^*_{s,t-1} + \beta \mu_s(\overline{U}^*_{:,t})\right) \tag{18}$$

$$\overline{U}^*_{s,t} = \mathrm{G}_{s,t}\left(\overline{U}^*_{1:S,1:T}\right) \tag{19}$$

for $s = 1, .., S$ and $t = 1, .., T-1$. Analogous arguments, using Eq. 11 and Eq. 13 instead of Eq. 12, show the same for $t = 1$ and $t = T$. □

**Lemma 2.** *For $0 < 2\alpha + \beta < 0.261$ and $\alpha > 0, \beta > 0$, the mapping* G *is a contraction mapping.*

*Proof.* The proof of Lemma 2 will use the Davis-Kahan Theorem. Given $\overline{U}_{1:S,1:T}$, $\overline{U}'_{1:S,1:T}$, let $\Sigma_{s,t}$ and $\Sigma'_{s,t}$ denotes the matrices

$$\Sigma_{s,1} = U_{s,1} + \alpha \overline{U}_{s,2} + \beta \mu_s\left(\overline{U}_{:,1}\right) \quad \text{and} \quad \Sigma'_{s,1} = U_{s,1} + \alpha \overline{U}'_{s,2} + \beta \mu_s\left(\overline{U}_{:,1}\right) \tag{20}$$

$$\Sigma_{s,t} = U_{s,t} + \alpha \overline{U}_{s,t-1} + \alpha \overline{U}_{s,t+1} + \beta \mu_s\left(\overline{U}_{:,t}\right) \quad \text{and} \quad \Sigma'_{s,t} = U_{s,t} + \alpha \overline{U}'_{s,t-1} + \alpha \overline{U}'_{s,t+1} + \beta \mu_s\left(\overline{U}_{:,t}\right) \tag{21}$$

$$\Sigma_{s,T} = U_{s,T} + \alpha \overline{U}_{s,T-1} + \beta \mu_s\left(\overline{U}_{:,T}\right) \quad \text{and} \quad \Sigma'_{s,T} = U_{s,T} + \alpha \overline{U}'_{s,T-1} + \beta \mu_s\left(\overline{U}_{:,T}\right) \tag{22}$$

so that $\mathrm{G}_{s,t}(\overline{U}_{1:S,1:T}) = \Pi_K(\Sigma_{s,t})$. Let $V_{s,t}$ and $V'_{s,t}$ denote orthonormal basis for the first $K$ eigenvectors of $\Sigma_{s,t}$ and $\Sigma'_{s,t}$, respectively. The following chain of equations can be seen to hold for $t = 2, .., T-1$, and $s = 1, .., S$:

$$\left\|\mathrm{G}_{s,t}(\overline{U}_{1:S,1:T}) - \mathrm{G}_{s,t}(\overline{U}'_{1:S,1:T})\right\|_F = \left\|\Pi_K(\Sigma_{s,t}) - \Pi_K(\Sigma'_{s,t})\right\|_F \tag{23}$$

$$= \left\|V_{s,t} R R^T V_{s,t}^T - V'_{s,t} V'^T_{s,t}\right\|_F \tag{24}$$

$$= \left\|\left(V_{s,t}R - V'_{s,t}\right) R^T V_{s,t}^T + V'_{s,t}\left(V_{s,t}R - V'_{s,t}\right)^T\right\|_F \tag{25}$$

$$\leq \left\|\left(V_{s,t}R - V'_{s,t}\right) R^T V_{s,t}^T\right\|_F + \left\|V'_{s,t}\left(V_{s,t}R - V'_{s,t}\right)^T\right\|_F \tag{26}$$

$$= \left\|\left(V_{s,t}R - V'_{s,t}\right) R^T\right\|_F + \left\|\left(V_{s,t}R - V'_{s,t}\right)^T\right\|_F \tag{27}$$

$$= 2\left\|\left(V_{s,t}R - V'_{s,t}\right)^T\right\|_F \tag{28}$$

$$\leq \frac{2\sqrt{2}}{\delta} \left\|\Sigma_{s,t} - \Sigma'_{s,t}\right\|_F \tag{29}$$

$$= \frac{2\sqrt{2}}{\delta} \left\|\alpha\left(\overline{U}_{s,t+1} - \overline{U}'_{s,t+1} + \overline{U}_{s,t-1} - \overline{U}'_{s,t-1}\right) + \beta\left(\mu_s\left(\overline{U}_{:,t}\right) - \mu_s\left(\overline{U}'_{:,t}\right)\right)\right\|_F \tag{30}$$

$$\leq \frac{2\sqrt{2}}{\delta} \left(\alpha\left(\left\|\overline{U}_{s,t+1} - \overline{U}'_{s,t+1}\right\|_F + \left\|\overline{U}_{s,t-1} - \overline{U}'_{s,t-1}\right\|_F\right) + \beta\left\|\mu_s\left(\overline{U}_{:,t}\right) - \mu_s\left(\overline{U}'_{:,t}\right)\right\|_F\right) \tag{31}$$

$$\leq \frac{2\sqrt{2}}{1 - 2\alpha - \beta} \left(\alpha\left(\left\|\overline{U}_{s,t+1} - \overline{U}'_{s,t+1}\right\|_F + \left\|\overline{U}_{s,t-1} - \overline{U}'_{s,t-1}\right\|_F\right) + \beta\left\|\mu_s\left(\overline{U}_{:,t}\right) - \mu_s\left(\overline{U}'_{:,t}\right)\right\|_F\right) \tag{32}$$

where we used the following steps for each step:

- Eq. 24: Let $R \in \mathbb{R}^{K \times K}$ satisfying $RR^T = I$. $R$ will satisfy additional condition of Davis-Kahan theorem.

†If $A$ and $B$ are positive semi-definite, then $\alpha A + \beta B$ is also positive semi-definite for non-negative $\alpha, \beta$.

- Eq. 25: Algebraic manipulation and re-arrangement.
- Eq. 26: Applying the triangle inequality.
- Eq. 27: Using the fact that $\left\|M^T V^T\right\|_F^2 = \mathrm{Tr}\left(M^T V^T V M\right) = \|M\|_F^2$ for any $V \in \mathcal{V}$.
- Eq. 28: Using the fact that $R$ is an orthogonal matrix and preserves Frobenius norm.
- Eq. 29: Using the Davis-Kahan theorem. $\delta$ is the difference between the $K$th and $(K+1)$th eigenvalues for $\Sigma_{s,t}$.
- Eq. 30: Algebraic manipulation and re-arrangement.
- Eq. 31: Applying the triangle inequality.
- Eq. 32: Given $U_{s,t}, \overline{U}_{s,t-1}, \overline{U}_{s,t+1}, \mu_s\left(\overline{U}_{:,t}\right)$, Weyl's inequality implies that the $K$th eigenvalue of $U_{s,t} + (\alpha\overline{U}_{s,t-1} + \alpha\overline{U}_{s,t+1} + \beta\mu_s\left(\overline{U}_{:,t}\right))$ is at least 1, and the $(K+1)$th eigenvalue is at most $2\alpha+\beta$, and hence, $\delta \geq 1 - 2\alpha - \beta$.

Analogous arguments are also valid for different time steps, and similar conditions also hold for $t = 1$ and $t =$, for $s = 1\,..\,S$:

$$\left\|\mathrm{G}_{s,1}\left(\overline{U}_{1:S,1:T}\right) - \mathrm{G}_{s,1}\left(\overline{U}'_{1:S,1:T}\right)\right\|_F \leq \frac{2\sqrt{2}}{1-2\alpha-\beta}\left(\alpha\left\|\overline{U}_{s,2} - \overline{U}'_{s,2}\right\|_F + \beta\left\|\left(\mu_s\left(\overline{U}_{:,1}\right) - \mu_s\left(\overline{U}'_{:,1}\right)\right)\right\|_F\right) \tag{33}$$

$$\left\|\mathrm{G}_{s,T}\left(\overline{U}_{1:S,1:T}\right) - \mathrm{G}_{s,T}\left(\overline{U}'_{1:S,1:T}\right)\right\|_F \leq \frac{2\sqrt{2}}{1-2\alpha-\beta}\left(\alpha\left\|\overline{U}_{s,T-1} - \overline{U}'_{s,T-1}\right\|_F + \beta\left\|\mu_s\left(\overline{U}_{:,T}\right) - \mu_s\left(\overline{U}'_{:,T}\right)\right\|_F\right) \tag{34}$$

$$\tag{35}$$

Our goal is to sum Eq. 33, Eq. 34 and Eq. 32 for each time step, and derive a general inequality. In order to show each step clearly, summation over terms with $\alpha$ and $\beta$ coefficient presented separately.

Summing terms with $\alpha$ coefficient (temporal smoothing), over each time step $t = 1,..,T$ and each subject $s = 1,..,S$ yields

$$\sum_{s=1}^{S}\left(\sum_{t=2}^{T-1}\alpha\left(\left\|\overline{U}_{s,t+1} - \overline{U}'_{s,t+1}\right\|_F + \left\|\overline{U}_{s,t-1} - \overline{U}'_{s,t-1}\right\|_F\right)\right) + \alpha\left(\left\|\overline{U}_{s,2} - \overline{U}'_{s,2}\right\|_F + \left\|\overline{U}_{s,T-1} - \overline{U}'_{s,T-1}\right\|_F\right) \tag{36}$$

$$= \sum_{s=1}^{S}\left(\sum_{t=2}^{T-1}2\alpha\left\|\overline{U}_{s,t} - \overline{U}'_{s,t}\right\|_F\right) + \alpha\left(\left\|\overline{U}_{s,1} - \overline{U}'_{s,1}\right\|_F + \left\|\overline{U}_{s,T} - \overline{U}'_{s,T}\right\|_F\right) \tag{37}$$

$$\leq \sum_{s=1}^{S}\sum_{t=1}^{T}2\alpha\left\|\overline{U}_{s,t} - \overline{U}'_{s,t}\right\|_F. \tag{38}$$

Similarly, summing terms with $\beta$ coefficient (subject-wise information sharing), over each time step and each subject yields:

$$\beta \sum_{s=1}^{S}\sum_{t=1}^{T} \left\|\mu_s\left(\overline{U}_{:,t}\right)-\mu_s\left(\overline{U}'_{:,t}\right)\right\|_F = \beta \sum_{s=1}^{S}\sum_{t=1}^{T} \left\| \frac{1}{S-1}\sum_{s^\dagger\in[K]\setminus s} \overline{U}_{s^\dagger,t} - \frac{1}{S-1}\sum_{s^\dagger\in[K]\setminus s} \overline{U}'_{s^\dagger,t} \right\|_F \tag{39}$$

$$= \beta \sum_{s=1}^{S}\sum_{t=1}^{T} \frac{1}{S-1} \left\| \sum_{s^\dagger\in[K]\setminus s} \left(\overline{U}_{s^\dagger,t} - \overline{U}'_{s^\dagger,t}\right) \right\|_F \tag{40}$$

$$\leq \beta \sum_{s=1}^{S}\sum_{t=1}^{T} \frac{1}{S-1} \sum_{s^\dagger\in[K]\setminus s} \left\| \overline{U}_{s^\dagger,t} - \overline{U}'_{s^\dagger,t} \right\|_F \tag{41}$$

$$= \beta \sum_{s=1}^{S}\sum_{t=1}^{T} \frac{1}{S-1} \left( \left( \sum_{s^\dagger=1}^{S} \left\| \overline{U}_{s^\dagger,t} - \overline{U}'_{s^\dagger,t} \right\|_F \right) - \left\| \overline{U}_{s,t} - \overline{U}'_{s,t} \right\|_F \right) \tag{42}$$

$$= \beta \sum_{t=1}^{T} \frac{1}{S-1} \sum_{s=1}^{S} (S-1) \left\| \overline{U}_{s,t} - \overline{U}'_{s,t} \right\|_F \tag{43}$$

$$= \beta \sum_{t=1}^{T}\sum_{s=1}^{S} \left\| \overline{U}_{s,t} - \overline{U}'_{s,t} \right\|_F . \tag{44}$$

Overall, combining $\alpha$ and $\beta$ terms, we end up with the following sum:

$$\sum_{t=1}^{T}\sum_{s=1}^{S} \beta \left\| \overline{U}_{s,t} - \overline{U}'_{s,t} \right\|_F + 2\alpha \left\| \overline{U}_{s,t} - \overline{U}'_{s,t} \right\|_F \tag{45}$$

$$= (\beta+2\alpha) \sum_{t=1}^{T}\sum_{s=1}^{S} \left\| \overline{U}_{s,t} - \overline{U}'_{s,t} \right\|_F . \tag{46}$$

Inserting the above summation results in the following inequality:

$$\sum_{t=1}^{T}\sum_{s=1}^{S} \left\| \mathrm{G}_{s,t}\left(\overline{U}_{1:S,1:T}\right) - \mathrm{G}_{s,t}\left(\overline{U}'_{1:S,1:T}\right) \right\|_F \leq \frac{2\sqrt{2}}{1-2\alpha-\beta}(2\alpha+\beta) \sum_{t=1}^{T}\sum_{s=1}^{S} \left\| \overline{U}_{s,t} - \overline{U}'_{s,t} \right\|_F \tag{47}$$

so that, when $0 \leq \frac{2\sqrt{2}}{1-2\alpha-\beta}(2\alpha+\beta) < 1$, G is a contraction mapping with the metric given below;

$$\mathrm{d}\left(\overline{U}_{1:S,1:T}, \overline{U}'_{1:S,1:T}\right) = \sum_{t=1}^{T}\sum_{s=1}^{S} \left\| \overline{U}_{s,t} - \overline{U}'_{s,t} \right\|_F . \tag{48}$$

As a conclusion, we have a bounded interval for $2\alpha+\beta$:

$$0 \leq 2\alpha+\beta < \frac{1}{1+2\sqrt{2}} \tag{49}$$

$$0 \leq 2\alpha+\beta < 0.261 \tag{50}$$

□

**Theorem 3** (Convergence of the Eigenvector Smoothing)**.**

$$\min_{\substack{\overline{U}_{s,t} \\ s=1,\ldots,S \\ t=1,\ldots,T}} \sum_{s=1}^{S} \left( \sum_{t=1}^{T} \left\|U_{s,t} - \overline{U}_{s,t}\right\|_F^2 + \sum_{t=1}^{T} \beta \left\|\overline{U}_{s,t} - \mu_s\left(\overline{U}_{:,t}\right)\right\|_F^2 + \sum_{t=1}^{T-1} \alpha \left\|\overline{U}_{s,t} - \overline{U}_{s,t+1}\right\|_F^2 \right) \tag{51}$$

subject to $\overline{U}_{s,t} \in \left\{VV^T \ : \ V \in \mathbb{R}^{G\times K},\ V^TV = I\right\} \ \forall s, \forall t.$

$$
\begin{aligned}
\overline{U}_{s,1}^{\ell+1} &= \Pi_K \left(U_{s,1} + \alpha \overline{U}_{s,2}^{\ell} + \beta \mu_s \left(\overline{U}_{:,1}^{\ell}\right)\right) \\
\overline{U}_{s,T}^{\ell+1} &= \Pi_K \left(\alpha \overline{U}_{s,T-1}^{\ell} + U_{s,T} + \beta \mu_s \left(\overline{U}_{:,T}^{\ell}\right)\right) \\
\overline{U}_{s,t}^{\ell+1} &= \Pi_K \left(\alpha \overline{U}_{s,t-1}^{\ell} + U_{s,t} + \alpha \overline{U}_{s,t+1}^{\ell} + \beta \mu_s \left(\overline{U}_{:,t}^{\ell}\right)\right), \quad t = 2, \ldots, T-1 \\
\overline{U}_{st}^{0} &= U_{st}, \quad t = 1, \ldots, T,
\end{aligned}
\tag{52}
$$

*Proof.* Eq. 51 must have a global minimum, since its feasible region is bounded and its objective function is bounded from below by 0. By Lemma 1, the global minimum of Eq. 51 must be a fixed point of G. By Lemma 2, G is a contraction mapping and Contraction Mapping Theorem (Theorem 1 implies that it has a unique fixed point. It follows that the unique fixed point of G is the global minimum of Eq. 51.

Since the Eigenvector Smoothing iterations are $\overline{U}_{1:S,1:T}^{\ell+1} = \mathrm{G}\left(\overline{U}_{1:S,1:T}^{\ell}\right)$, Lemma 2 and Theorem 1 imply that $\overline{U}_{1:S,1:T}^{\ell+1}$ converges to the fixed point of G, as $l \to \infty$. Thus $\overline{U}_{1:S,1:T}^{\ell+1}$ converges to the global minimum of Eq. 51, proving the theorem. □

#### S2.2 Convergence in Simulation Experiments

Fig. S1 illustrates the convergence of MuDCoD with different $\alpha$ and $\beta$ values by quantifying the $\mathrm{L}_{1,1}$ norm of the difference of the estimated eigenvectors between consecutive iterations.

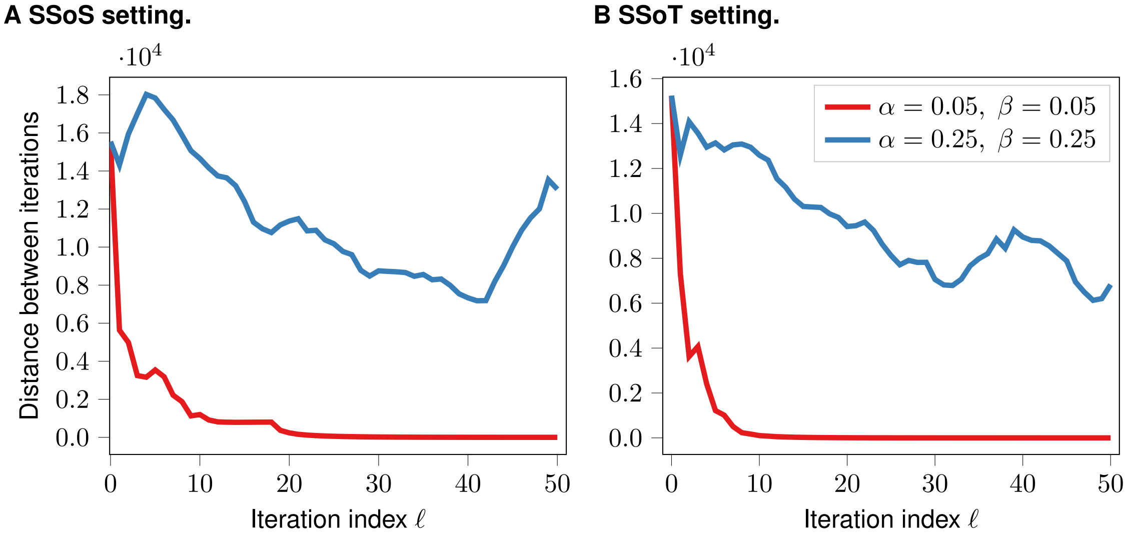


Figure S1: *Convergence status of MuDCoD with different $\alpha$ and $\beta$ values.* $y$-axis quantifies the $\mathrm{L}_{1,1}$ norm of the difference of estimated eigenvectors between consecutive iterations: $\sum_{s=1}^{S} \sum_{t=1}^{T} \|\overline{U}_{s,t}^{\ell} - \overline{U}_{s,t}^{\ell-1}\|_{1,1}$. Other parameters of the MuS-Dynamic-DCBM were set as follows: $G = 500$, $K = 10$, $p_{\text{in}} = (0.3, 0.3)$, $p_{\text{out}} = (0.1, 0.1)$, $r_{\text{time}} = 0.2$, $r_{\text{subject}} = 0.2$, $S = 4$, $T = 4$.

### S3 Further Implementation Details of the Methods

#### S3.1 Details of Betzel *et al.* [3]

Betzel-2019 [3] is a multi-layer modularity maximization method that has been developed within the context of brain networks. It is applicable to similar settings with multiple subjects or time points by treating connectivity matrices (e.g., co-expression matrices in our setting) as "layers" from multiple subjects or time points. Betzel-2019 operates on these multi-layer networks to detect modular structures across individuals and time points.

We obtained the Matlab implementation for Betzel-2019 from `https://www.brainnetworkslab.com/coderesources` and its required Matlab toolbox GenLouvain [7][13]. Betzel-2019 implementation has two free hyperparameters [3]. The structural resolution parameter, $\gamma$, modulates the sizes of the communities: smaller $\gamma$ leads to larger communities. The inter-subject coupling parameter, $\omega$, affects the consistency of communities across the layers (subjects/times): larger $\omega$ promotes information sharing across layers. Betzel-2019 provides an efficient parameter sampling approach that surveys a large portion of the two dimensional hyperparameter space. In our application of Betzel-2019, we used their default hyperparameter sampling approach to narrow down the hyperparameter space, and then followed their method to generate communities at different combinations of these two hyperparameters. Due to the variability caused by Betzel-2019's sensitivity to hyperparameters, we summarized the results by considering parameter values that resulted in the most frequent number of detected communities. More specifically, we partitioned the numbers of communities detected under different hyperparameter settings into several intervals, and used the subset of hyperparameters with results in the highest frequency interval. Communities from each of these hyperparameter settings were evaluated, and mean performance metrics were reported for each simulation replicate.

#### S3.2 Details of Liu *et al.* [10]

The Original MATLAB implementation of PisCES [9] is publicly available at `https://github.com/letitiaLiu/PisCES`. We re-implemented PisCES in Python together with MuDCoD and made it available at `https://github.com/bo1929/MuDCoD`. We choose the smoothing hyperparameter $\alpha$ and the number of communities $K$ with the same procedures described in Liu *et al.* [9].

### S4 Related Literature

| Reference | Multi-subject | Multi-time | Method |
|---|---|---|---|
| Norman and Cicek [14] | Y | Y | Spatio-temporal Steiner tree |
| Bassett *et al.* [1] | N | Y | Modularity maximization |
| Betzel-2019 [2] | Y | Y | Multi-layer modularity maximization |
| Ma *et al.* [11] | N | Y | Graph regularized evolutionary NMF |
| Cribben and Yu [5] | Y | Y | Network change point detection |
| Matias and Miele [12] | N | Y | Dynamic SBM |
| Xu and Hero [18] | N | Y | Dynamic SBM |
| Ting *et al.* [17] | Y | Y | Multi-subject & Markov-switching SBM |
| Chi *et al.* [4] | N | Y | Smoothed spectral clustering |
| PisCES [9] | N | Y | Smoothed spectral clustering |
| MuDCoD | Y | Y | Smoothed spectral clustering |

Table S1: Summary of recent network community analysis methods relevant for multi-subjects and/or dynamic nature. Column "Multi-subject" ("Multi-time") indicates if the study directly mentions and analyses multi-subject (or multi-time points) networks: Y (yes) or N (no). In the column "Method", the abbreviations are as follows: NMF: non-negative matrix factorization, SBM: stochastic block model.

#### S5 Empirical Motivation for MuDCoD's Formulation

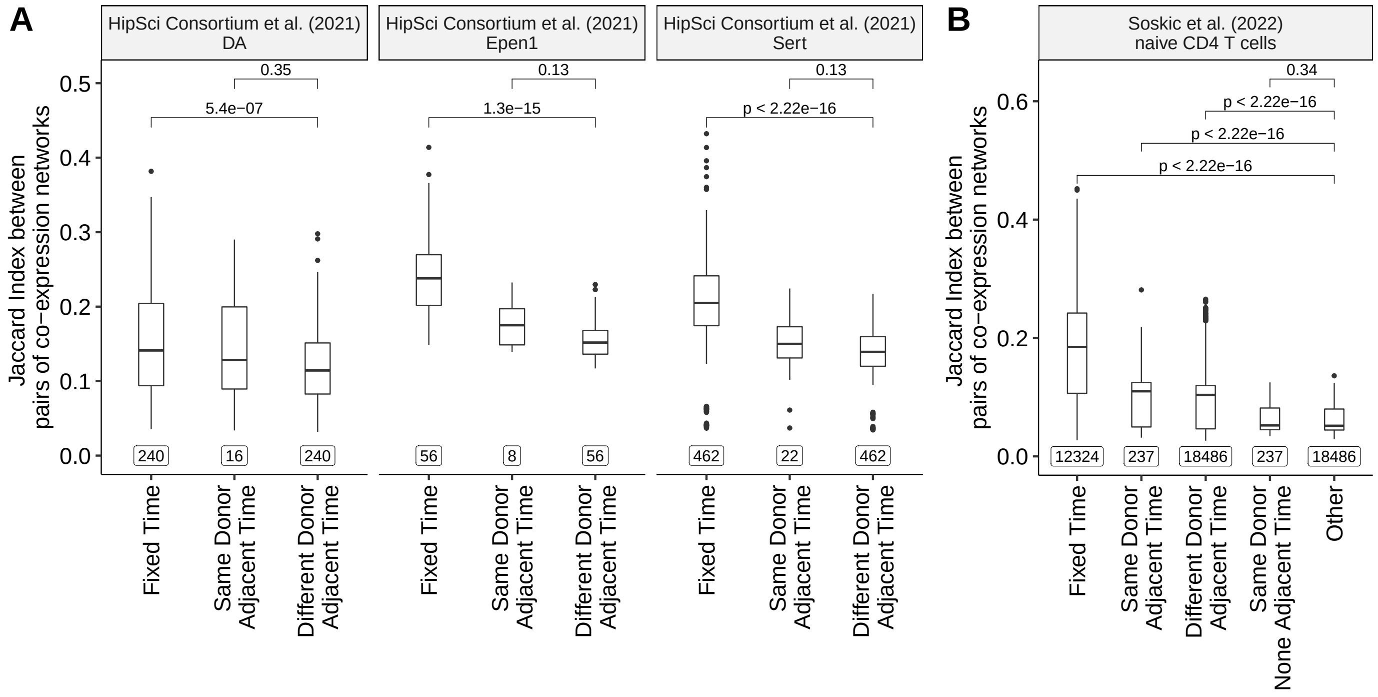


Figure S2: Comparison of the input donor and time-specific adjacency matrices (co-expression networks, $A_{s,t}$, $s = 1 \cdots, S$, $t = 1, \cdots, T$ ). A. For cell types, DA Epen1, and Sert of the HipSci Consortium *et al.* [6] scRNA-seq dataset. B. For naive human CD4$^+$ T cells of the Soskic *et al.* [16] scRNA-seq dataset. For all the cell types analyzed, donor adjacency matrices are compared across the following groups: 1. *Fixed time*: Pairwise comparison of donor adjacency matrices at fixed time point ($A_{s_1,t}$ vs. $A_{s_2,t}$, where $s_1, s_2 \in \{1, \cdots, S\}$, $s_1 \neq s_2$, $t = 1, \cdots, T$); *Same Donor/Adjacent Time*: Comparison of individual donor's adjacency matrices from adjacent time points ($A_{s,t_1}$ vs. $A_{s,t_2}$, where $s = 1, \cdots, S$, $t_1 = 1, \cdots, T-1$, $t_2 = t_1 + 1$); *Different Donor/Adjacent Time*: Comparison of adjacency matrices of two donors at adjacent time points ($A_{s_1,t_1}$ vs. $A_{s_2,t_2}$, where $s_1, s_2 \in \{1, \cdots, S\}$, $s_1 \neq s_2$, $t_1 = 1, \cdots, T-1$, $t_2 = t_1 + 1$); *Same Donor/Nonadjacent Time*: Comparison of individual donor's adjacency matrices from nonadjacent time points ($A_{s,t_1}$ vs. $A_{s,t_2}$, where $s = 1, \cdots, S$, $t_1 = 1, \cdots, T$, $t_2 \neq t_1 + 1$, $t_2 \neq t_1 - 1$), only applicable when there are more than two time points; *Other*: Rest of pairwise comparisons. Higher Jaccard index for "Fixed Time" compared to "Other" or "Different Donor/Adjacent Time" motivates smoothness among spectral representations of the subjects at the same time point. Similarly, higher Jaccard index for "Same Donor/Adjacent Time" motivates smoothness of spectral representations over the adjacent time points. Numbers below each boxplot represent the numbers of comparisons.

### S6 Analysis and Interpretation of the Inferred Communities

#### S6.1 Summary of Communities Inferred by MuDCoD on the scRNA-seq Datasets

Table S2: Summaries of MuDCoD inferred communities for each cell type and time point across the two scRNA-seq dataset applications. The first rows of each dataset report the mean and standard deviation of the total number of communities across donors. The second and third rows report the median, 10-th, and 90-th percentiles of the community sizes and densities, respectively.

| **HipSci Consortium *et al.* [6]** | **DA** | | **Sert** | | **Epen1** | |
|---|---|---|---|---|---|---|
| | Day-30 | Day-52 | Day-30 | Day-52 | Day-30 | Day-52 |
| **Number of communities** mean $\pm$ SD | $9.25 \pm 2.2$ | $14.6 \pm 3.7$ | $15.7 \pm 2.7$ | $16.4 \pm 3.8$ | $16.5 \pm 3.9$ | $17.8 \pm 4.2$ |
| **Size of comm.** $P_{10\%}$/median/$P_{90\%}$ | 51/143/458 | 25/95/293 | 41/98/280 | 3/96/291 | 3/125/259 | 3/91/306 |
| **Density of comm.** $P_{10\%}$/median/$P_{90\%}$ | .03/.30/.46 | .04/.35/.51 | .03/.38/.55 | .02/.39/.55 | .02/.39/.59 | .01/.39/.64 |

| **Soskic *et al.* [16]** | **Naive CD4$^+$ T** | | | |
|---|---|---|---|---|
| | 0-hours | 16-hours | 40-hours | 5-days |
| **Number of communities** mean $\pm$ SD | $6.34 \mp 3.51$ | $6.8 \mp 3.42$ | $6.3 \mp 3.11$ | $6.41 \mp 3.69$ |
| **Size of comm.** $P_{10\%}$/median/$P_{90\%}$ | 3/19/242 | 3/17/219 | 3/17/220 | 3/19/222 |
| **Density of comm.** $P_{10\%}$/median/$P_{90\%}$ | .04/.07/.25 | .04/.08/.29 | .05/.08/.22 | .04/.07/.21 |

##### S6.1.1 Details of HipSci Consortium *et al.* [6]

Table S2 reports characteristics of MuDCoD inferred gene modules across each time point for each cell type. While the numbers of donors available for each cell type were different, the numbers of inferred communities, with the exception of DA at day-30, were similar in size. The mean number of communities across donors ranges between 9.25 and 17.8. The low standard deviation in the number of communities indicates that the number of communities across donors does not drastically change. We next computed the density of the communities by dividing the number of co-expression edges by the total number of possible edges. The median community densities range between 0.30 to 0.39 for all cell types. DA at day-30 leads to the sparsest communities with larger community sizes. The median density of the communities at day-52 is either the same or larger than the communities discovered at day-30.

##### S6.1.2 Details of Soskic *et al.* [16]

Table S2 reports characteristics of MuDCoD inferred gene modules at each time point and reveals that the median number of communities which vary between 6.3 and 6.8 across time points is remarkably stable. The relatively high standard deviation of the median number of communities, i.e., in the range of 3.11 to 3.51, elucidate that the number of communities across donors tends to exhibit marked changes. The inferred gene modules are also highly similar in size for all-time points, with a median size within the range $[17, 19]$, and with a notable exception of few relatively larger communities for the time point 0-hours (90-th percentile size of 242).

#### S6.2 Gene Set Enrichment Analysis of the Communities Inferred by MuDCoD for Individual Donors Across the Time Points in the HipSci Consortium *et al.*, (2021) dataset

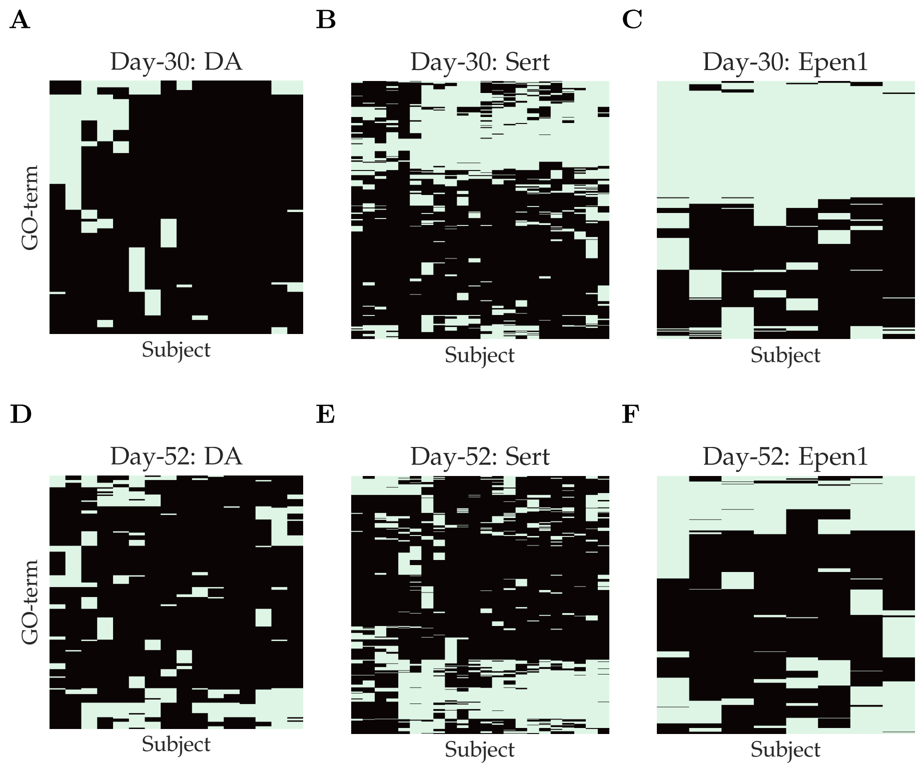


Figure S3: Gene set enrichment analysis using GO biological processes was carried out for each of the inferred communities (with a size of at least 20 genes) at each time point and for each donor with `clusterProfiler`. Background gene sets were limited to those used in the construction of the adjacency matrices, and FDR was controlled at level 0.05 with the Benjamini-Hochberg procedure. Enriched GO terms (rows) of the donors (columns) are depicted with the green color. Heatmaps highlight the overall patterns of enriched GO terms across the subjects for each "time point-cell type" combination. The GO term names on the rows are suppressed to improve visualization.

#### S6.3 NMI Scores Between Inferred Communities of Donor Pairs for Soskic *et al.*, (2022) dataset

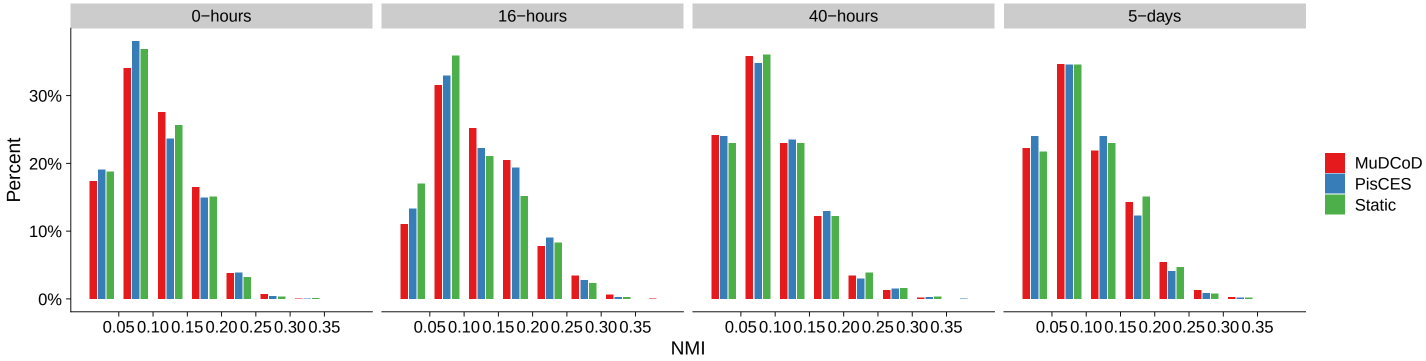


Figure S4: NMI scores between inferred communities of donor pairs for CD4$^+$ T cells at each time point. NMI scores between gene communities of every pair of donors at 0-hours, and 16-hours, 40-hours, and 5-days after stimulation, respectively in first, second, third and fourth panels. The $y$-axis denotes the percentage of donor pairs. Both $x$-axis and $y$-axis are in logarithmic scale.

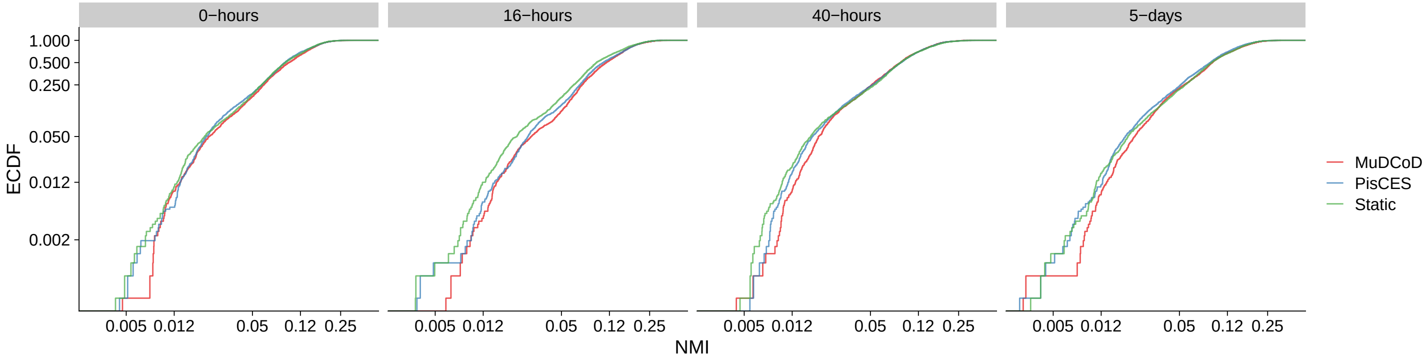


Figure S5: Empirical cumulative distribution functions (ECDF) of NMI scores between inferred communities of donor pairs for CD4$^+$ T cells at each time point. NMI scores between gene communities of every pair of donors at 0-hours, and 16-hours, 40-hours, and 5-days after stimulation. The $y$-axis denotes the cumulative proportion of donor pairs below the corresponding NMI score at the $x$-axis.

#### S6.4 NMI Scores Between Inferred Communities of Donor Pairs for HipSci Consortium *et al.*, (2021) dataset

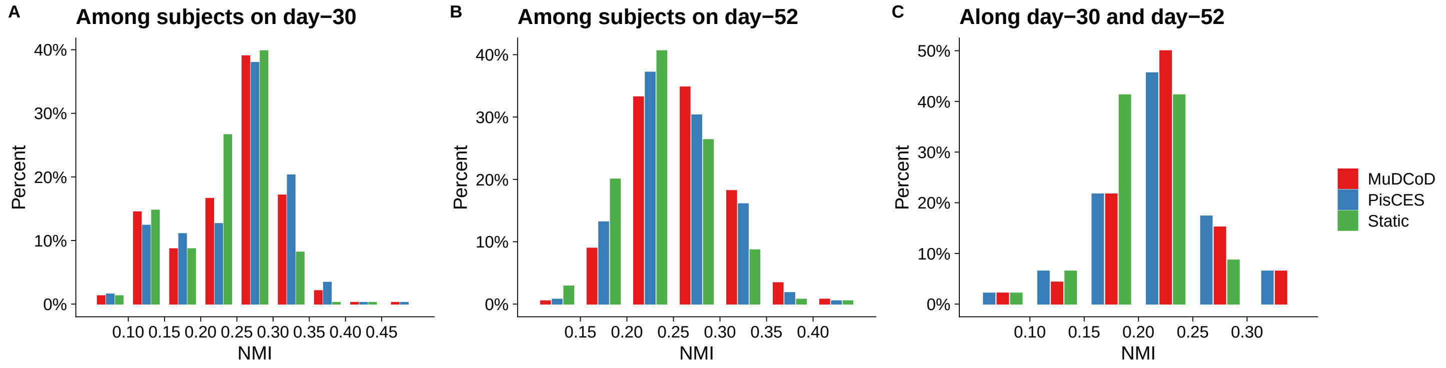


Figure S6: NMI scores between inferred gene modules of donor and time point pairs aggregated across all cell types. (A) and (B) quantify NMI scores between modules of every pair of donors on day-30 and day-52, respectively. (C) displays NMI scores between inferred gene modules of each donor on day-30 and on day-52. The $y$-axis denotes the percentage of donor pairs. Both $x$-axis and $y$-axis are in logarithmic scale.

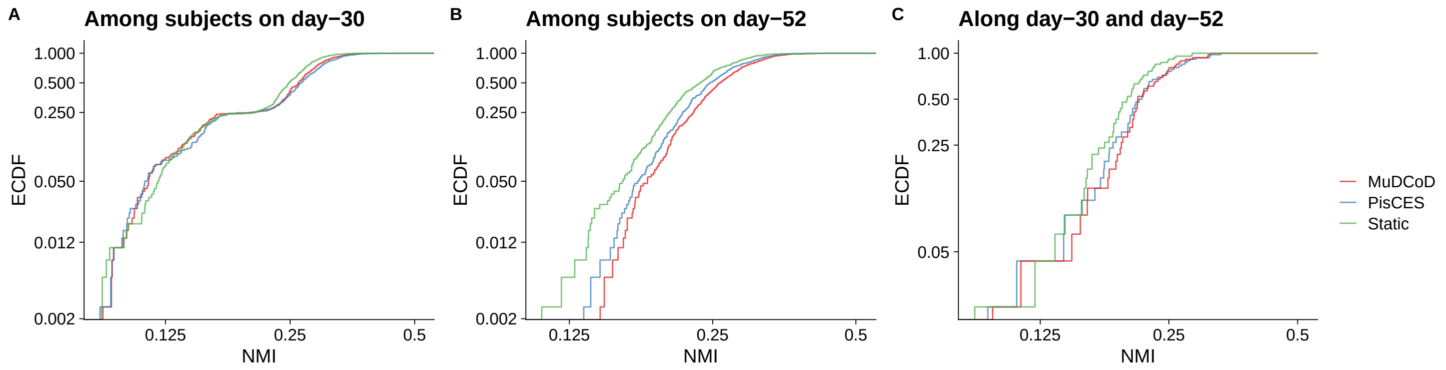


Figure S7: Empirical cumulative distribution functions (ECDF) of NMI scores between inferred gene modules of donor and time point pairs aggregated across all cell types. (A) and (B) quantify NMI scores between modules of every pair of donors on day-30 and day-52, respectively. (C) displays NMI scores between inferred gene modules of each donor on day-30 and on day-52. The $y$-axis denotes the cumulative proportion of donor pairs below the corresponding NMI score at the $x$-axis.

#### S6.5 Analysis of Communities Inferred by PisCES in HipSci Consortium *et al.*, (2021) dataset

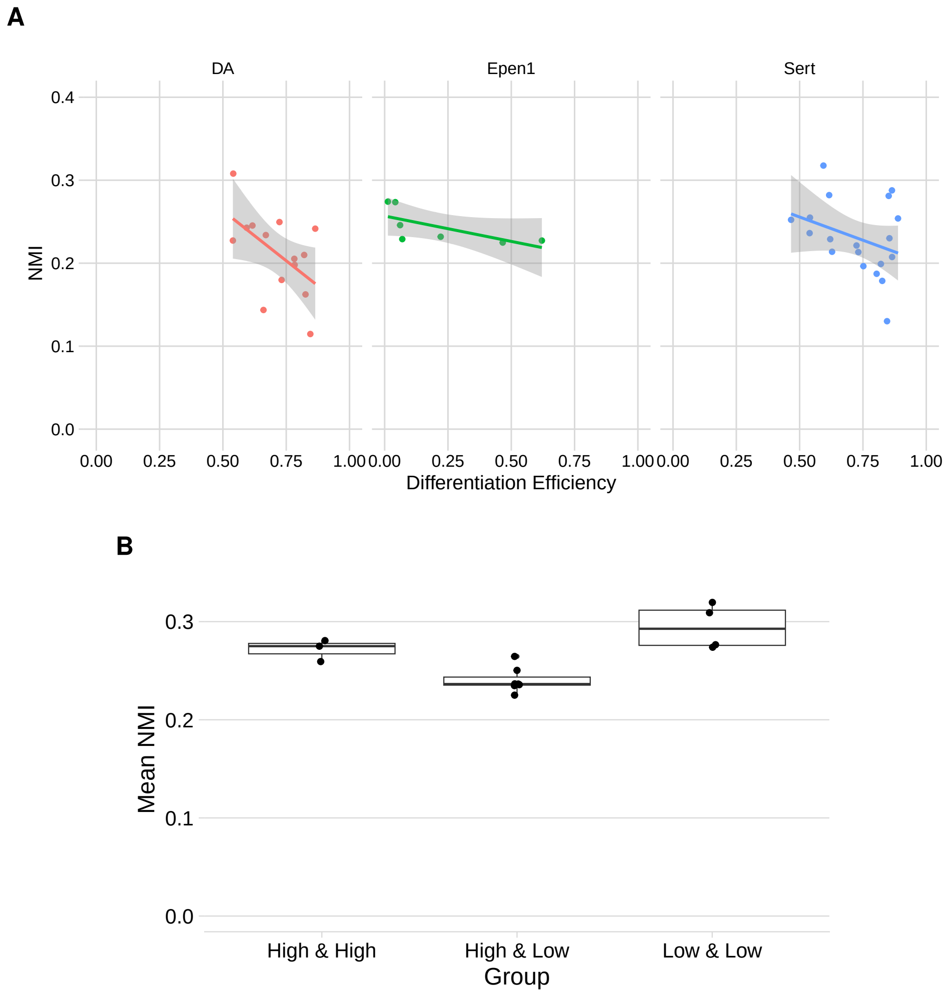


Figure S8: (A) NMI scores of each donor between the modules inferred by PisCES on day-30 and on day-52 against the differentiation efficiency. For each cell type (DA, Sert, Epen1), each donor's modules from day-30 and day-52 were compared with NMI and plotted against donor's differentiation efficiency. (B) Comparison of the mean NMI scores within and between the low and high differentiation efficiency groups of Epen1 cells. Donor labels for differentiation efficiency were obtained from [6]. NMI scores between pairs of donors are calculated based on their PisCES inferred modules. Differentiation efficiency groups were generated based on the percentiles of differentiation efficiency values across the donors, i.e., 50-th percentile (inclusive) corresponds to the low differentiation efficiency group.

#### S6.6 Analysis of Communities Inferred by Static Spectral Clustering in HipSci Consortium *et al.*, (2021) dataset

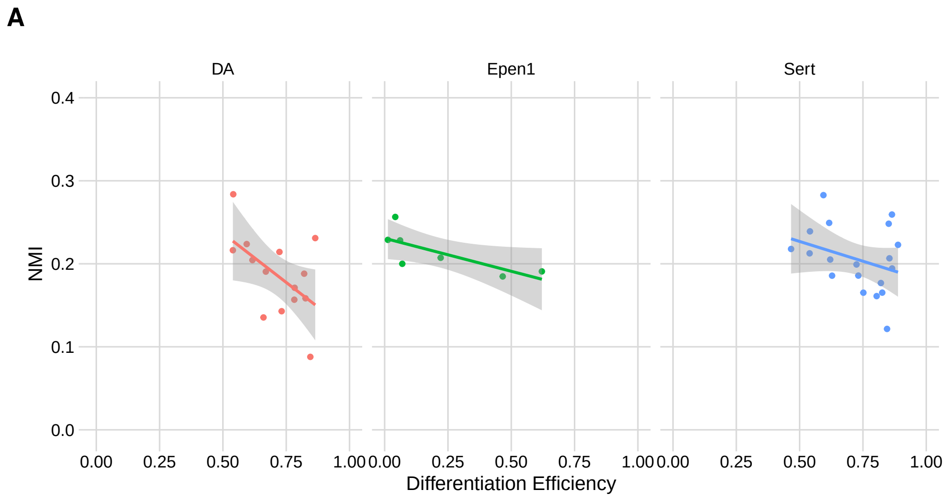


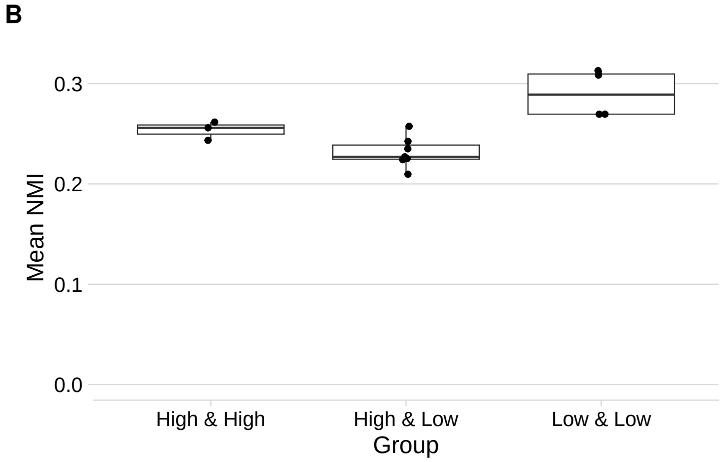


Figure S9: (A) NMI scores of each donor between the modules inferred by static spectral clustering on day-30 and on day-52 against the differentiation efficiency. For each cell type (DA, Sert, Epen1), each donor's modules from day-30 and day-52 were compared with NMI and plotted against donor's differentiation efficiency. (B) Comparison of the mean NMI scores within and between the low and high differentiation efficiency groups of Epen1 cells. Donor labels for differentiation efficiency were obtained from [6]. NMI scores between pairs of donors are calculated based on their static spectral clustering inferred modules. Differentiation efficiency groups were generated based on the percentiles of differentiation efficiency values across the donors, i.e., 50-th percentile (inclusive) corresponds to the low differentiation efficiency group.

#### S6.7 Analysis of Communities Inferred by PisCES in Soskic *et al.*, (2022) dataset

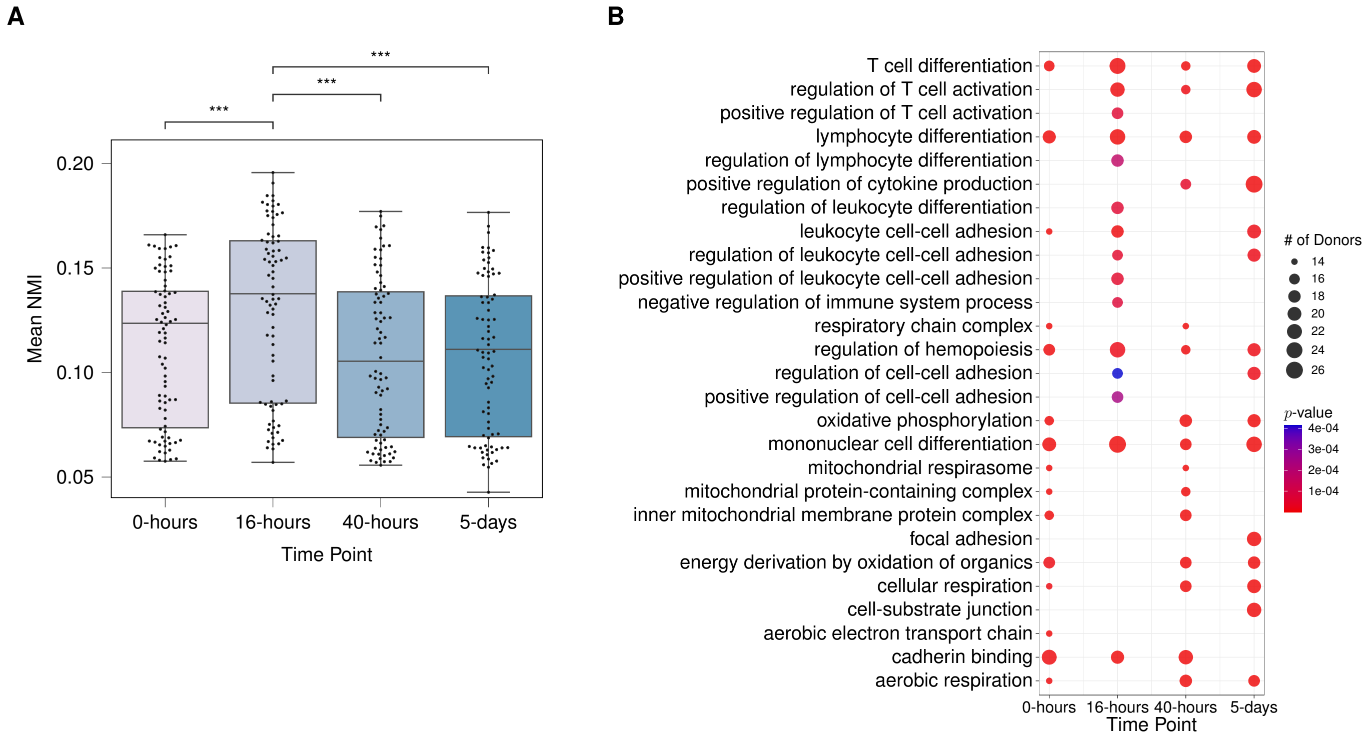


Figure S10: (A) Normalized mutual information scores between pairs of donors based on their gene modules inferred by PisCES at each time point. Each data point stands for a donor, and the $y$-axis denotes the mean of NMI scores between that donor and other donors at the corresponding time point. All the statistically significant comparisons with time point 16-hours from the one-sided Wilcoxon rank-sum test are marked (with $p\text{-value}\leq 0.05$), *** stands for $p\text{-value}\leq 0.001$. (B) Set of top fifteen frequent significantly enriched biological processes (with the adjusted $p$-values $\leq 0.05$) of most prioritized communities contributed by each donor at different time points. Overall, $79 \times 4 = 316$ gene sets were contributed by $79$ donors at $4$ time points, and correspondingly, $316$ separate enrichment analyses were performed. Displayed are significant biological processes, their corresponding number of appearances in the most prioritized communities of donors, and the minimum $p$-values of enrichment among those communities.

#### S6.8 Analysis of Communities Inferred by Static Spectral Clustering in Soskic *et al.*, (2022) dataset

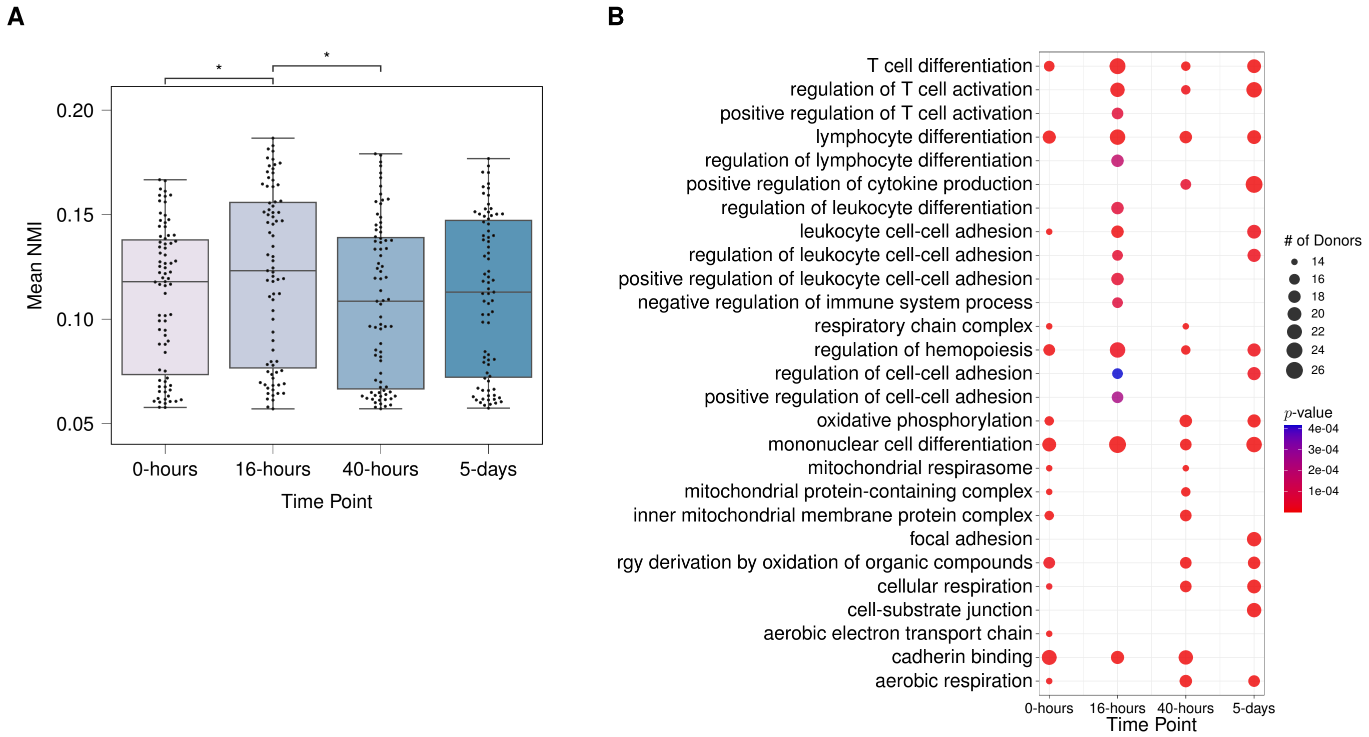


Figure S11: (A) Normalized mutual information scores between pairs of donors based on their gene communities inferred by static spectral clustering at each time point. Each data point stands for a donor, and the $y$-axis denotes the mean of NMI scores between that donor and other donors at the corresponding time point. All the statistically significant comparisons with time point 16-hours from the one-sided Wilcoxon rank-sum test are marked (with $p$-value$\leq 0.05$), $\star$ stands for $0.01 < p\text{-value} \leq 0.05$. (B) Set of top fifteen frequent significantly enriched biological processes (with the adjusted $p$-values $\leq 0.05$) of most prioritized communities contributed by each donor at different time points. Overall, $79 \times 4 = 316$ gene sets were contributed by $79$ donors at $4$ time points, and correspondingly, $316$ separate enrichment analyses were performed. Displayed are significant biological processes, their corresponding number of appearances in the most prioritized communities of donors, and the minimum $p$-values of enrichment among those communities.

### S7 Runtime Comparisons

All analyses on real datasets were carried out on a single node of a cluster with an Intel Xeon E5-2640 CPU (2.40GHz). On the HipSci Consortium *et al.* [6] dataset, the typical runtimes of MuDCoD and PisCES were comparable. For three different cell types, runtimes ranged between 30 minutes to 1.5 hours for about $2,000$ genes. Similarly, when applied to the Soskic *et al.* [16] dataset, the runtimes of MuDCoD and PisCES were similar, typically falling within the 20 minutes to 1 hour range.

Fig. S12 presents a detailed comparison of the runtimes of PisCES and MuDCoD on simulated data on a laptop with Intel Core i5-10210U CPU (1.60GHz). We assessed the runtime requirements for community inference with fixed hyperparameters and hyperparameter tuning separately. When the hyperparameters are fixed, the inference times are very close and PisCES is slightly faster (Fig. S12A). However, MuDCoD requires much longer runtimes for the cross-validation of hyperparameters (Fig. S12B). This is mainly because we perform a grid search for two hyperparameters ($\alpha$ and $\beta$) opposed to a single parameter ($\alpha$) for PisCES.

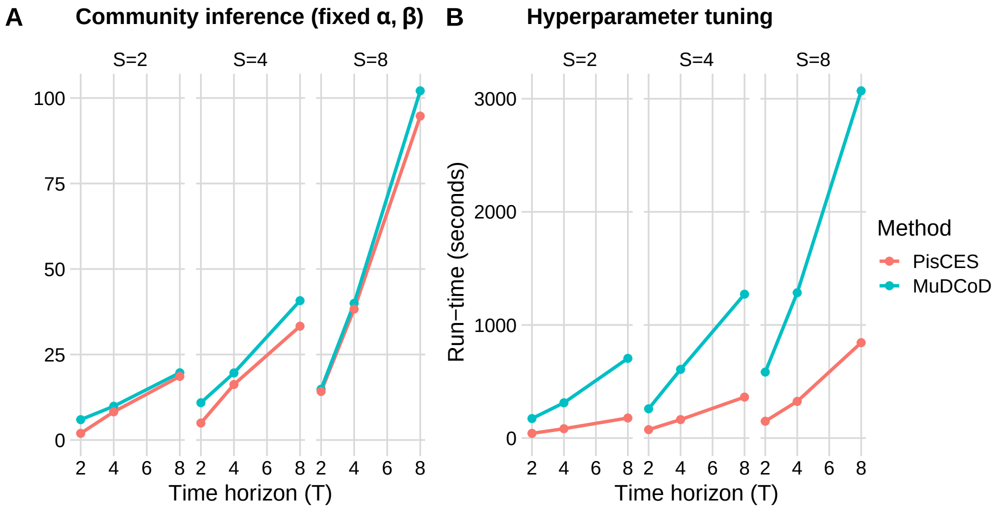


Figure S12: Runtime comparisons of MuDCoD and PisCES community inference with fixed hyperparameters (A) and hyperparameter tuning (B) on simulated data. The simulation parameters were set as follows for the SSoS setting: the network size $G$=200, the number of class labels $K$=4, the in-cluster and out-cluster density parameters $p_{\text{in}}$=$(0.2, 0.25)$ and $p_{\text{out}}$=$(0.1, 0.125)$, number of subjects $S \in \{2, 4, 8\}$, the number of time points $T \in \{2, 4, 8\}$, $r_{\text{subject}}$=0.35, and $r_{\text{time}}$=0.35. For cross-validation, we performed grid search for $\alpha \in \{0.01, 0.05, 0.1\}$ and $\beta \in \{0.01, 0.05, 0.1\}$.

### S8 ECDFs of absolute values of gene-gene correlations for the Soskic *et al.*, (2022) dataset

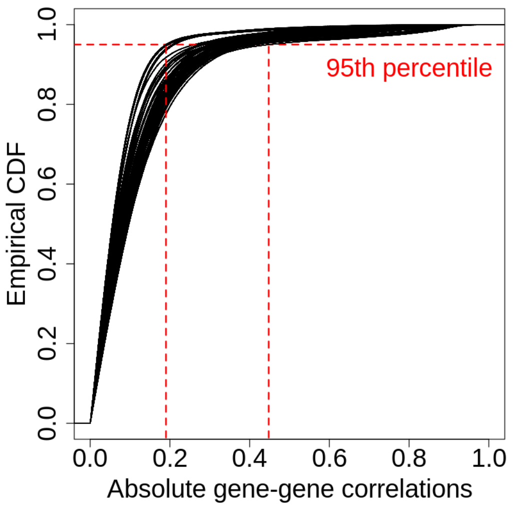


Figure S13: Empirical cumulative distribution functions (CDFs) of absolute values of the gene-gene correlations for 50 randomly sampled donors of the Soskic *et al.*, (2022). The vertical red lines depict the lowest and highest 95th percentiles of the absolute values of the gene-gene correlations across the donors.

#### References


[1] Bassett, D. S. *et al.* (2013). Robust detection of dynamic community structure in networks. *Chaos: An Interdisciplinary Journal of Nonlinear Science*, **23**(1), 013142.

[2] Betzel, R. F. *et al.* (2019a). The community structure of functional brain networks exhibits scale-specific patterns of inter- and intra-subject variability. *NeuroImage*, **202**, 115990.

[3] Betzel, R. F. *et al.* (2019b). The community structure of functional brain networks exhibits scale-specific patterns of inter-and intra-subject variability. *Neuroimage*, **202**, 115990.

[4] Chi, Y. *et al.* (2007). Evolutionary spectral clustering by incorporating temporal smoothness. In *Proceedings of the 13th ACM SIGKDD international conference on Knowledge discovery and data mining*, pages 153–162.

[5] Cribben, I. and Yu, Y. (2017). Estimating whole-brain dynamics by using spectral clustering. *Journal of the Royal Statistical Society: Series C (Applied Statistics)*, **66**(3), 607–627.

[6] HipSci Consortium *et al.* (2021). Population-scale single-cell RNA-seq profiling across dopaminergic neuron differentiation. *Nature Genetics*, **53**(3), 304–312.

[7] Jutla, I. S. *et al.* (2011). A generalized louvain method for community detection implemented in matlab. *URL http://netwiki. amath. unc. edu/GenLouvain.*

[8] Li, T. *et al.* (2020). Network cross-validation by edge sampling. *Biometrika*, **107**(2), 257–276.

[9] Liu, F. *et al.* (2018a). Global spectral clustering in dynamic networks. *Proceedings of the National Academy of Sciences*, **115**(5), 927–932.

[10] Liu, F. *et al.* (2018b). Global spectral clustering in dynamic networks. *Proceedings of the National Academy of Sciences*, **115**(5), 927–932.

[11] Ma, X. *et al.* (2019). Detecting evolving communities in dynamic networks using graph regularized evolutionary nonnegative matrix factorization. *Physica A: Statistical Mechanics and its Applications*, **530**, 121279.

[12] Matias, C. and Miele, V. (2017). Statistical clustering of temporal networks through a dynamic stochastic block model. *Journal of the Royal Statistical Society: Series B*, **79**(4), 1119–1141.

[13] Mucha, P. J. *et al.* (2010). Community structure in time-dependent, multiscale, and multiplex networks. *science*, **328**(5980), 876–878.

[14] Norman, U. and Cicek, A. E. (2019). ST-Steiner: a spatio-temporal gene discovery algorithm. *Bioinformatics*, **35**(18), 3433–3440.

[15] Shen, H.-W. and Cheng, X.-Q. (2010). Spectral methods for the detection of network community structure: a comparative analysis. *Journal of Statistical Mechanics: Theory and Experiment*, **2010**(10), P10020.

[16] Soskic, B. *et al.* (2022). Immune disease risk variants regulate gene expression dynamics during CD4+ T cell activation. *Nature Genetics*, **54**(6), 817–826. Number: 6 Publisher: Nature Publishing Group.

[17] Ting, C.-M. *et al.* (2021). Detecting Dynamic Community Structure in Functional Brain Networks Across Individuals: A Multilayer Approach. *IEEE Transactions on Medical Imaging*, **40**(2), 468–480. Conference Name: IEEE Transactions on Medical Imaging.

[18] Xu, K. S. and Hero, A. O. (2014). Dynamic stochastic blockmodels for time-evolving social networks. *IEEE Journal of Selected Topics in Signal Processing*, **8**(4), 552–562.
